# Different mutant RUNX1 oncoproteins program alternate haematopoietic differentiation trajectories

**DOI:** 10.1101/2020.08.10.244657

**Authors:** Sophie G Kellaway, Peter Keane, Benjamin Edginton-White, Regha Kakkad, Ella Kennett, Constanze Bonifer

## Abstract

Mutations of the hematopoietic master regulator RUNX1 cause acute myeloid leukaemia, familial platelet disorder and other haematological malignancies whose phenotypes and prognoses depend upon the class of RUNX1 mutation. The biochemical behaviour of these oncoproteins and their ability to cause unique diseases has been well studied, but the genomic basis of their differential action is unknown. To address this question we compared integrated phenotypic, transcriptomic and genomic data from cells expressing four types of RUNX1 oncoproteins in an inducible fashion during blood development from embryonic stem cells. We show that each class of mutated RUNX1 deregulates endogenous RUNX1 function by a different mechanism, leading to specific alterations in developmentally controlled transcription factor binding and chromatin programming. The result is distinct perturbations in the trajectories of gene regulatory network changes underlying blood cell development that are consistent with the nature of the final disease phenotype. The development of novel treatments for RUNX1-driven diseases will therefore require individual consideration.

## Introduction

RUNX1 is a transcription factor which is absolutely essential for hematopoietic development both in vivo and in vitro (Lacaud *et al*, 2002; Okuda *et al*, 1996). In humans, different classes of RUNX1 mutations lead to distinct malignancies and clinical outcomes (Bellissimo & Speck, 2017). Mutations involving RUNX1 are one of the most common recurrent drivers of acute myeloid leukaemia (AML) found in around 14% of cases (Papaemmanuil *et al*, 2016), but also cause other haematological conditions such as familial platelet disorder (FPD) as well as acute lymphoblastic leukaemia (ALL) (Schlegelberger & Heller, 2017), and are associated with chronic myelogenous leukaemia (CML) (Lugthart *et al*, 2010). Established leukemic cells carrying different types of RUNX1 mutations show specific transcriptional and chromatin profiles (Assi *et al*, 2019). However, in patients, RUNX1 mutations are associated with additional genetic alterations that disrupt differentiation and alter cellular growth (Gaidzik *et al*, 2016). Therefore, the molecular mechanisms how the sole expression of different types of RUNX1-oncoproteins drive the development of specific disease phenotypes is unclear.

RUNX1 mutations can occur affecting the DNA-binding domain (DBD), transactivation domain (TAD) or are a result of translocations resulting in the generation of fusion proteins. RUNX1 functions by directly binding DNA together with its obligate partner CBFβ via the DBD, in large complexes mediated by the TAD (Koh *et al*, 2013; Petrovick *et al*, 1998; Wotton *et al*, 1994). After hematopoietic stem cells (HSCs) have formed, its continued expression during differentiation is not essential but helps to correctly pattern and maintain cells in the correct lineage balance (Cai *et al*, 2011; Chen *et al*, 2009; Tober *et al*, 2013), in concert with other transcription factors such as the GATA, C/EBP and ETS families (Beck *et al*, 2013; Burda *et al*, 2010; Goode *et al*, 2016). Mutations in the DBD are typically point mutations which abrogate binding of RUNX1 to DNA but leave the rest of the protein intact, these are found as germline mutations in FPD but can also be seen in AML (Song *et al*, 1999). Premature stop codons or frameshift mutations typically remove the TAD, but may or may not affect the DBD. These are typically found in AML with poor prognosis (Döhner *et al*, 2017; Gaidzik *et al*, 2016; Mendler *et al*, 2012) but have also been seen in FPD (Song *et al.*, 1999). Recurrent translocations include t(8;21), t(3;21) and t(12;21), and these result in the fusion of part of the RUNX1 protein to all or part of another protein – ETO, EVI1 and ETV6 in the examples given – and are found in AML, CML and ALL (Golub *et al*, 1995; Mitani *et al*, 1994; Miyoshi *et al*, 1993; Romana *et al*, 1995).

The biochemical properties of mutant RUNX1 proteins are well characterised. DBD mutated proteins, as expected, cannot bind DNA; they have limited nuclear localisation but maintain CBFβ interaction (Matheny *et al*, 2007; Michaud *et al*, 2002). TAD mutants can bind DNA with varying efficiency and maintain CBFβ interactions, but show very limited nuclear localisation (Matheny *et al*, 2007; Michaud *et al*, 2002). Fusion proteins which maintain the RUNX1 DBD are still able to bind DNA, but further interactions are translocation specific – for example RUNX1-ETO interacts with repressive complexes (Amann *et al*, 2001). When deleted in HSCs of mice, straightforward RUNX1 deficiency causes an increase in immature myeloid cell formation, thrombocytopenia and lymphocytopenia (Putz *et al*, 2006; Sun & Downing, 2004). Expression of RUNX1 DBD mutated proteins in mice induce more complex phenotypes including myelodysplasia, mixed lineage cells and a reduction in colony forming progenitor cells in the aorta/gonad/mesonephros (Cammenga *et al*, 2007; Matheny *et al*, 2007; Watanabe-Okochi *et al*, 2008). TAD mutant proteins on the other hand, show dosage dependent phenotypes in mice, with severe disruption to formation of blood across multiple lineages (Matheny *et al*, 2007; Watanabe-Okochi *et al*, 2008). Similarly, expression of fusion proteins such as RUNX1-ETO and RUNX1-EVI1 in mice leads to large scale disruption of blood formation from haematopoietic progenitors with increased self-renewal (Maki *et al*, 2005; Okuda *et al*, 1998).

It is unclear precisely how RUNX1 mutant proteins contribute to disease. Initial hypotheses that these mutations lead to haploinsufficiency of RUNX1 or mediate dominant negative effects do not fully explain disease phenotypes (Cai *et al*, 2000; Cammenga *et al*, 2007; Matheny *et al*, 2007). We therefore carried out a parallel comparative study on two RUNX1 mutants representing DBD and TAD mutations, and two RUNX1 translocations and investigated how they affect transcriptional control and RUNX1 driven gene regulatory networks in haematopoietic progenitors. We show that each RUNX1 mutant protein interferes with the RUNX1 driven gene regulatory network in its own way, setting up distinct chromatin landscapes and leading to divergent outcomes of progenitor development.

## Results

### Mutant RUNX1 proteins disrupt haematopoietic differentiation

To understand the individual action of mutant RUNX1 proteins, we utilised a well characterised embryonic stem cell differentiation system which recapitulates the different steps of haematopoietic specification of blood cells from haemogenic endothelium, and allows inducible expression of oncoproteins (Goode *et al*, 2016; Iacovino *et al*, 2011; Lancrin *et al*, 2010; Regha *et al*, 2015). We induced each mutant with doxycycline (dox) in otherwise healthy blood progenitor cells (progenitors) at the onset of the RUNX1 transcriptional program (Figure 1A). The RUNX1 mutations studied were R201Q, also reported as R174Q dependent on the RUNX1 isoform, which is a DBD mutant, R204X (also reported as R177X) which is truncated following the DBD, and the fusion proteins RUNX1-ETO and RUNX1-EVI1 (Figure 1A). Induction conditions of each mutant RUNX1 protein were adjusted such that expression levels were approximately equal to the wild type RUNX1 (Supplementary Figure 1A, Kellaway *et al*, 2020; Regha *et al*, 2015). As differentiation in this system is not entirely synchronous, timing of induction was adjusted in a cell line specific manner such that it occurred in approximately the same target cell populations ensuring that results were comparable (Supplementary Figure 1B).

**Figure 1 -.**
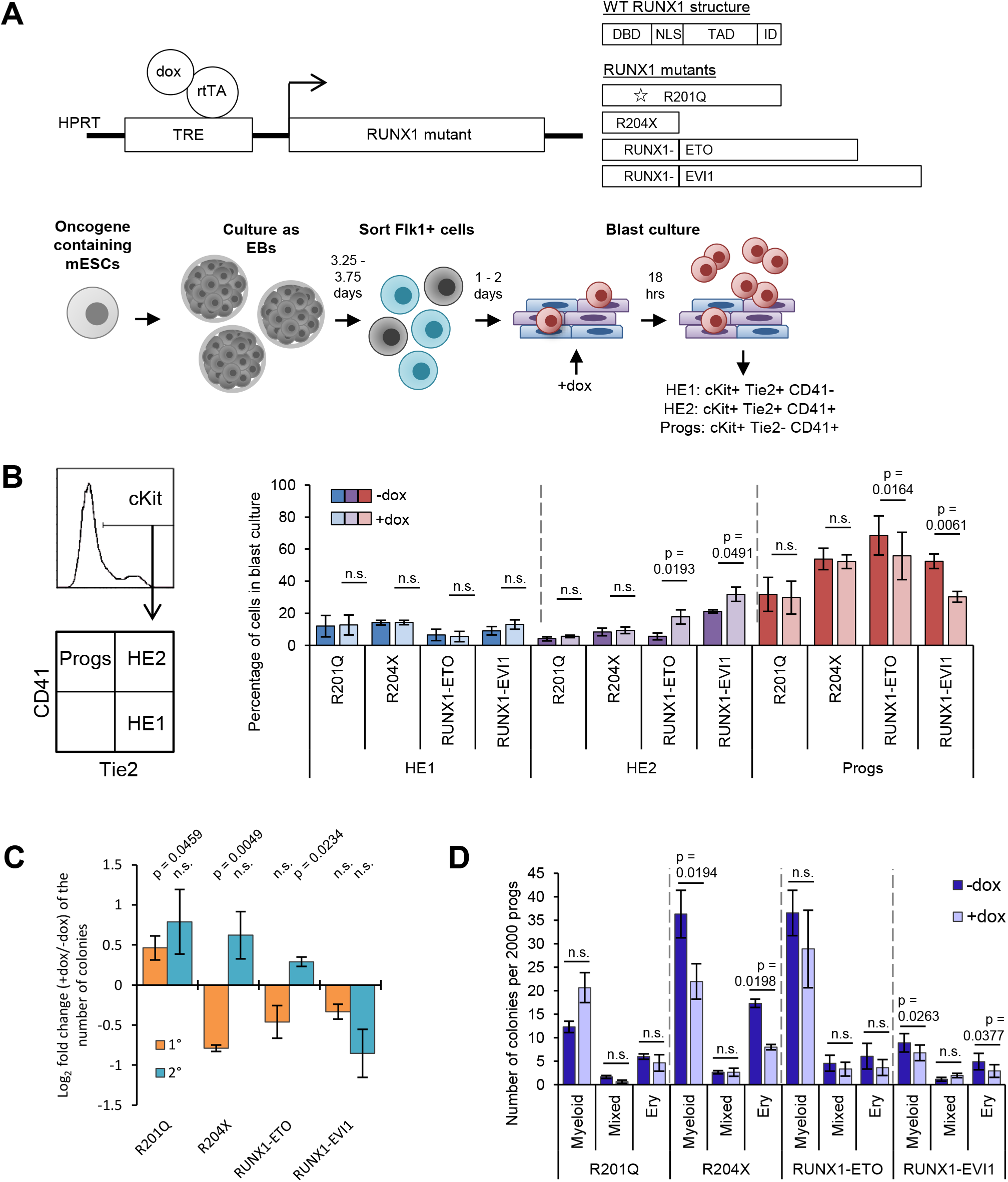
Induction of RUNX1 mutants during blood differentiation perturbs progenitor identity. A Schematic showing the RUNX1 inducible constructs used, the embryonic stem cell differentiation system and the stage of induction of the transgenes. B Flow cytometry was used to assess the proportion of cells in the blast culture which were HE1, HE2 or progenitors as indicated in the schematic on the left. Bars show the mean percentage of cells in each population. N = 3 for R201Q, n = 4 for R204X and RUNX1-ETO and n = 5 for RUNX1-EVI1. C Progenitors were placed into colony forming assays in the absence of continued dox. The bars show log2 fold change of induced (+dox) by non-induced (-dox) for primary colonies in orange, and secondary colonies in blue. R201Q primary colony forming n = 5, n = 3 for all others. D The absolute number of colonies of each lineage subtype from the primary colony forming assays in (C) is shown. Data information: (B-D) error bars show standard error of the mean. P-values were calculated using paired t-tests between – and +dox pairs, n.s. indicates p > 0.05.

We first assessed the impact of the RUNX1 mutant proteins on hematopoietic development. We have previously shown that RUNX1-ETO and RUNX1-EVI1 impede the endothelial-haematopoietic transition (EHT) for which RUNX1 is required (Kellaway *et al*, 2020; Regha *et al*, 2015), causing a reduced proportion of progenitors and increased proportion of late haemogenic endothelium (HE2) cells, indicating that fusion proteins were acting as dominant negative to the endogenous RUNX1. In contrast, no effect on the EHT was observed with either R201Q or R204X (Figure 1B, Supplementary Figure 1C). We next investigated how each RUNX1 mutation affected terminal differentiation and self-renewal ability of haematopoietic progenitors. In serial replating assays we found that the RUNX1 mutants behaved in a disease-specific fashion (Figure 1C-D). R201Q caused an increase in clonogenicity in both primary and secondary colony forming assays. In addition, fewer megakaryocytes were observed to form following induction of R201Q in the mixed lineage colonies (Supplementary Figure 1D). Expression of R204X and RUNX1-ETO which both cause AML led to an initial reduction in clonogenicity across all lineages, but an increase upon replating, indicative of a differentiation block and enhanced self-renewal. RUNX1-EVI1 caused a reduction in both primary and secondary colony forming capacity, again across all lineages, presumably due to the lineage decision promiscuity and cell cycle defects we have previously observed for this protein (Kellaway *et al*, 2020).

In summary, the four RUNX1 oncoproteins disrupt terminal differentiation in colony forming assays, reflecting the different diseases which they cause, but only the two translocations affected the RUNX1 dependent EHT.

### Endogenous RUNX1 binding changes in response to the presence of oncogenic RUNX1

To investigate the molecular basis of the observed phenotypic differences, we performed RNA-seq, ATAC-seq and ChIP-seq experiments in c-Kit+ progenitors (Supplementary Figure 2A) following induction of each of the mutant forms of RUNX1 and integrated the data. We found that changes to chromatin accessibility and gene expression were largely driven by mutant-specific changes in the endogenous RUNX1 binding patterns (Figure 2). R201Q triggered only minor changes to chromatin accessibility and gene expression following induction but caused a surprising large scale reduction in endogenous RUNX1 binding (Figure 2A, Supplementary Figure 2B). This reduction was not caused by direct competition with RUNX1 binding to chromatin as we were unable to detect binding of the R201Q protein by ChIP, using an antibody against the HA tag (Supplementary Figure 2C) and was reproducibly found in multiple manual ChIP experiments in cases where we obtained insufficient material to produce a sequencing library. In contrast, induction of the R204X protein caused little disruption to endogenous RUNX1 binding, but greater changes to chromatin accessibility, and again was not found to directly bind chromatin. We cannot exclude the possibility that R201Q and R204X can bind chromatin in a transient fashion, but the signal was below the detection limits of the ChIP experiments. Sites with altered chromatin accessibility following induction of RUNX1-ETO and RUNX1-EVI1 were also seen, with those sites lost associated with loss of endogenous RUNX1, and sites gained associated with gain of RUNX1, following RUNX1-ETO and RUNX1-EVI1 displacing some of the endogenous RUNX1 (Kellaway *et al*, 2020; Regha *et al*, 2015). Furthermore, following induction of RUNX1-EVI1 we found an increase in total binding of RUNX1 (Supplementary Figure 2B). Genome browser screenshots showing the changes in RUNX1 binding are shown in Figure 2B and Supplementary Figure 2D, these also show that residual RUNX1 binding is preserved at some sites following induction of R201Q but not all.

**Figure 2 -.**
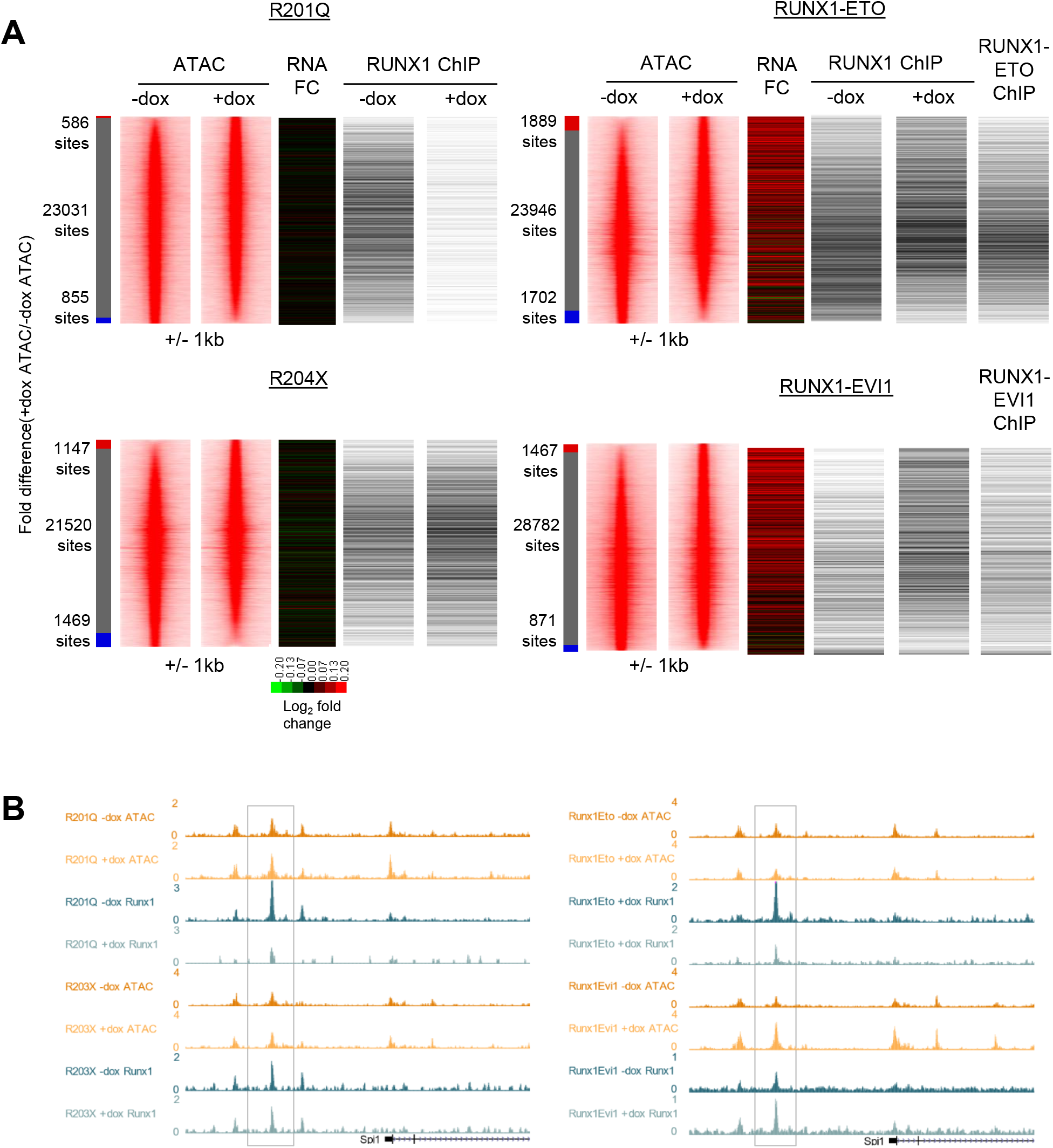
Mutant RUNX1 induction leads to specific changes to endogenous RUNX1 binding and chromatin accessibility. A Chromatin accessibility in cKit+CD41+Tie2-sorted progenitors at distal sites as determined by ATAC-seq was ranked by fold change of the +dox/-dox tag count and represented as density plots (+/- 1kb from the summit). The gene expression fold change as determined by RNA-seq (+dox/-dox) was plotted alongside based on nearest gene assigned. The binary presence or absence of a RUNX1, RUNX1-ETO or RUNX1-EVI1 ChIP peak was also plotted based on intersection with the open chromatin. The red bar indicates +dox specific sites, grey shared and blue -dox specific sites where the normalised tag-count of specific sites was at least 2-fold different. B UCSC Genome browser screenshot of CPM-normalised ATAC-seq and ChIP-seq tracks at the Spi1 locus. The box highlights the Spi1 enhancer which demonstrates changes in RUNX1 binding and chromatin accessibility.

### RUNX1 binding in the presence of mutant proteins is influenced by altered CBFβ interactions

We questioned whether the changes to endogenous RUNX1 may be due to the mutant proteins interfering with the binding of endogenous RUNX1 to CBFβ using in situ proximity ligation assays (PLA). By using antibodies specific to either the wild type RUNX1, HA-tagged induced mutant proteins or untagged RUNX1-EVI1, we assessed in single cells whether the induced RUNX1 oncoproteins were complexed with CBFβ, and quantified whether the interaction between CBFβ and endogenous RUNX1 was affected by oncoprotein induction. We first examined the localisation of the mutant RUNX1 proteins. Both RUNX1-ETO and RUNX1-EVI1 were clearly localised in nuclei (Figure 3A and Supplementary Figure 3A, left panels) whereas both R201Q and R204X exhibited diffuse staining with little protein found in the nucleus, consistent with previous studies (Michaud *et al*, 2002; Osato *et al*, 1999). We then examined whether induced proteins interacted with CBFβ and where. We found a high number of interactions between RUNX1-ETO and CBFβ, and RUNX1-EVI1 and CBFβ located within the nucleus (Figure 3A and Supplementary Figure 3A, right panels). In contrast, we observed very few interactions between R201Q and CBFβ, or R204X and CBFβ compared to background. Interestingly, despite minimal nuclear localised R201Q and R204X protein, we saw PLA foci in the nucleus, suggesting that some mutant RUNX1-containing complexes were capable of nuclear translocation.

**Figure 3 -.**
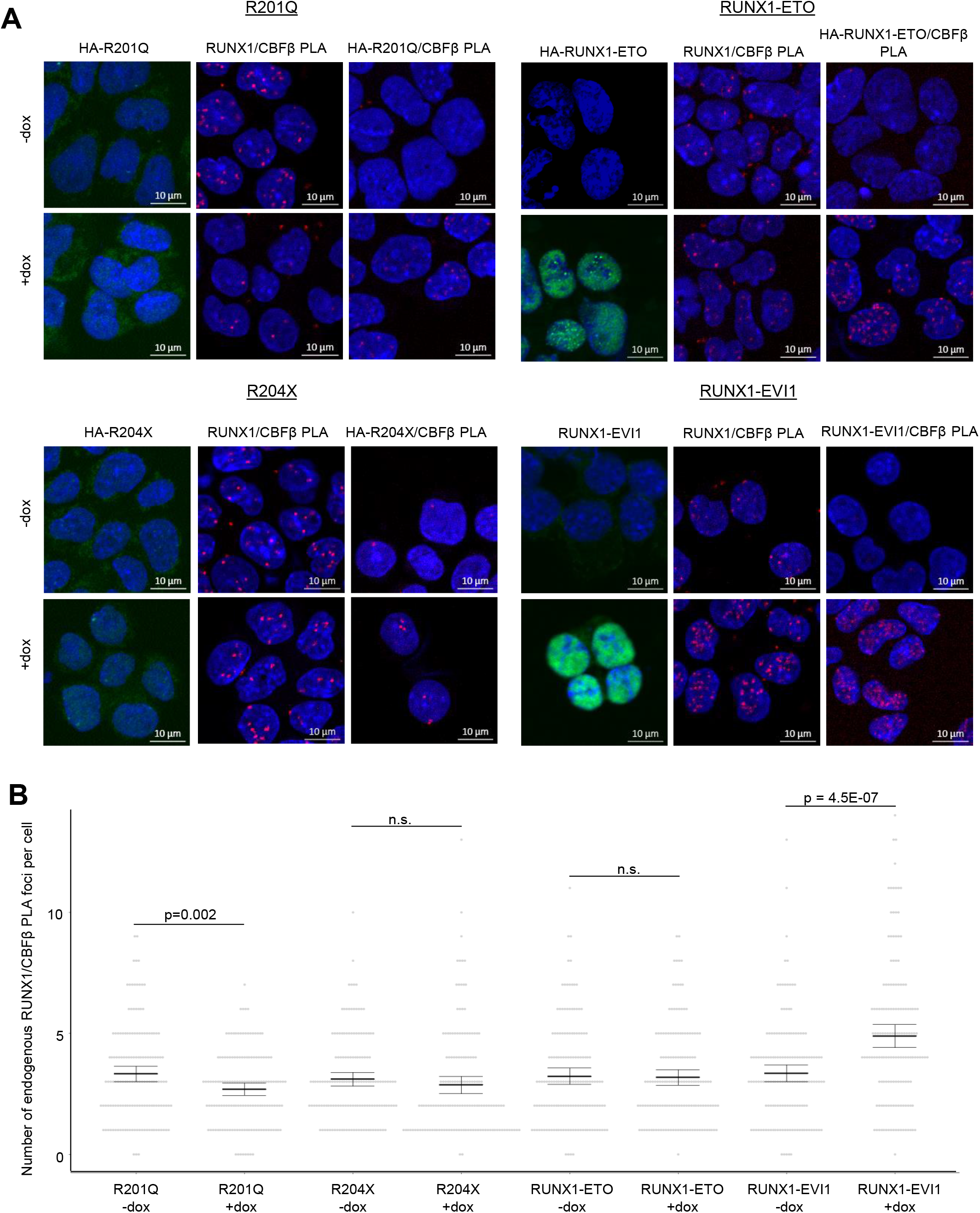
RUNX1 mutants interact with CBFβ and partially disrupt RUNX1/CBFβ interactions. A Representative images are shown of immunocytochemistry and PLA in progenitors with and without induction of the mutant forms of RUNX1. For each cell line, on the left is immunocytochemistry of the mutant protein alone (shown in green, using anti-HA or anti-EVI1 antibodies) counterstained with DAPI (blue). In the centre is PLA of endogenous RUNX1 with CBFβ (red), with DAPI (blue). On the right is PLA of the mutant RUNX1 with CBFβ (red), with DAPI (blue). B The number of endogenous RUNX1/CBFβ PLA foci were counted in 150 cells across 3 biological replicates and shown by the grey circles. The mean and 95% confidence intervals are indicated by the bar and error bar. P-values were calculated using two-sample T-tests between – and + dox pairs, n.s. indicates a p-value > 0.05.

We next assessed the quantity of interactions of the endogenous RUNX1 and CBFβ and compared them to the ChIP-seq results. Antibodies against wild type RUNX1 and CBFβ alone showed expected staining patterns which were unaffected by dox induction (Supplementary Figure 3B). In the uninduced cells, the number of PLA foci was similar for all cell lines allowing us to see only the effects of the mutant proteins (Figure 3B, p-value=0.723 by one way ANOVA). RUNX1-ETO and R204X expression caused no change to the quantity of RUNX1/CBFβ interactions, suggesting the changes in RUNX1 binding were due to displacement, as the endogenous RUNX1 protein was able to form complexes with CBFβ as normal. RUNX1-EVI1 caused an increase in the number of RUNX1/CBFβ foci (Figure 3A and Supplementary Figure 3A, centre panels) which mirrored the ChIP-seq data wherein we saw increased RUNX1 binding (Figure 2). Most strikingly however, given the mild phenotype, R201Q expression caused a reduction in the number of PLA foci, explaining the decrease in the amount of RUNX1 available to efficiently bind DNA as seen with the ChIP-seq result (Figure 2). The RUNX1 antibody used for this assay was unable to discriminate the endogenous RUNX1 from the induced R201Q and therefore some of these foci may in fact be R201Q/CBFβ interactions, meaning RUNX1/CBFβ interactions were even further reduced than measured.

Taken together, these data show that binding of the endogenous RUNX1 is disrupted by concurrent expression of mutant RUNX1 proteins, with concomitant variation in the frequency of interactions between endogenous RUNX1 and CBFβ. There was no evidence that CBFβ was stably sequestered by mutant RUNX1 proteins, although it is possible that CBFβ is sequestered and then degraded. Displacement of endogenous RUNX1 binding by the mutant RUNX1 proteins was only found in the case of the two fusion proteins.

### Changes in RUNX1 binding lead to mutation class-specific changes in gene regulation

To understand how the changes to the RUNX1 program drive the phenotypes observed and to see whether the mutant RUNX1 forms target similar transcriptional networks we compared gene expression changes. Overall gene expression data for both the induced and the uninduced state were highly consistent across the four cell lines (Supplementary Figure 4A) and replicates correlated well (Supplementary Figure 4B). As expected from the cell biological data, RUNX1-ETO and RUNX1-EVI1 de-regulated the most genes across the EHT, and similarly, fewer changes were seen following induction of R201Q and R204X (Figure 4A, Supplementary Figure 4C). With induction of R201Q the vast majority of genes continued to be regulated according to their expected trajectory, with a subset failing to be up-regulated to the extent they normally would including *Hba-a1, Cd79b* and *Mef2c.* The induction of RUNX1-ETO caused the greatest number of genes to not be down-regulated sufficiently including *Gfi1* ((Lancrin *et al*, 2012), Figure 4A). Looking specifically at the changes at the specific cell stages, RUNX1-ETO and RUNX1-EVI1 both caused the greatest number of genes to be up or downregulated in both HE2 and progenitors, and R204X only caused upregulation of genes at the HE2 stage and not in progenitors, for example *Mecom* and *Plek* (Supplementary Figure 4C).

**Figure 4 -.**
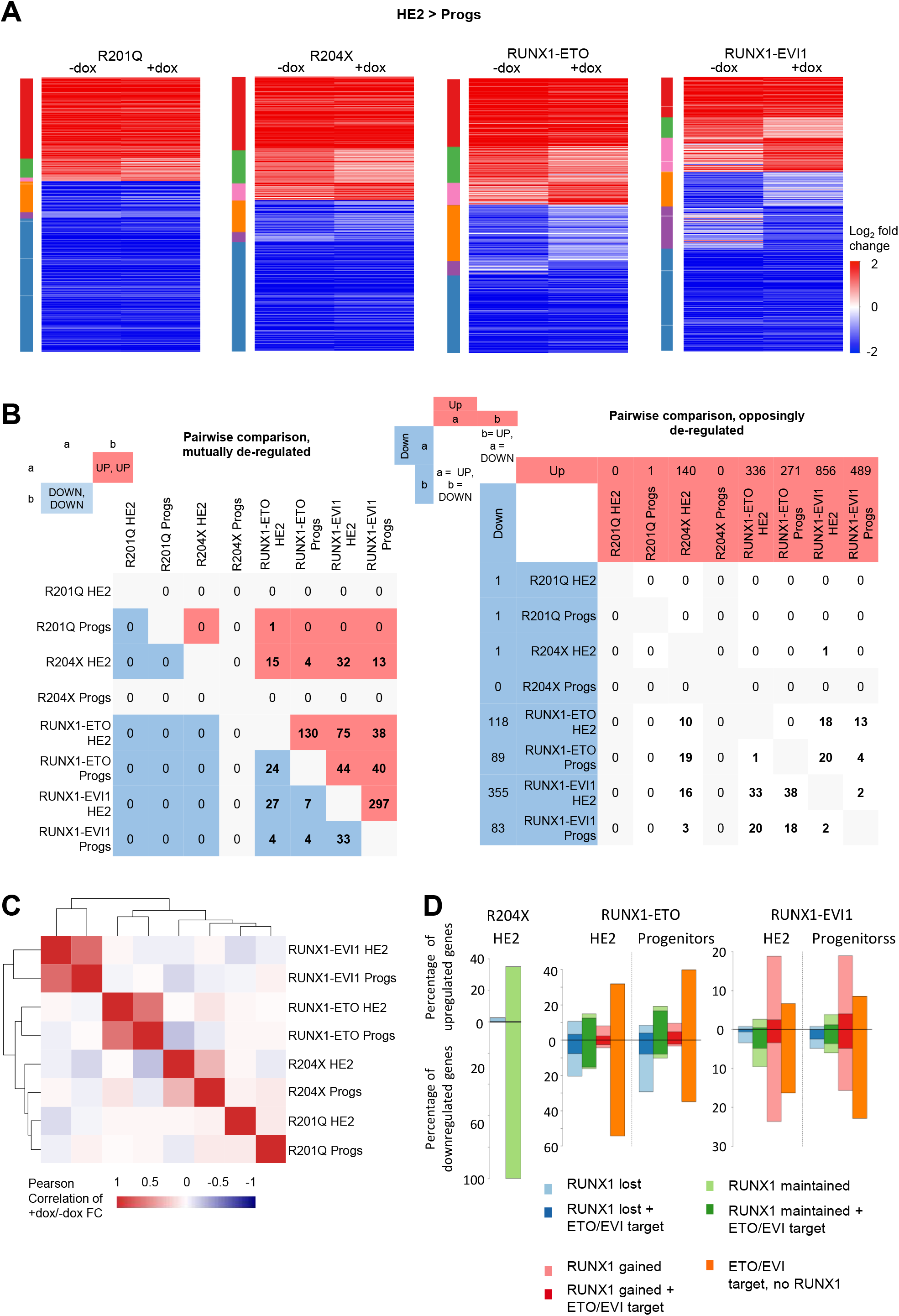
Mutant forms of RUNX1 cause unique and shared gene expression changes. A Heatmaps showing the log_2_ fold gene expression changes across the HE2 to progenitor transition. Colour bars on the left indicate genes which are (red) upregulated in both - and +dox, (green) upregulated in -dox only, (pink) upregulated in +dox only, (orange) downregulated in -dox only, (purple) downregulated in +dox only, (blue) downregulated in both - and +dox. B Pairwise analysis of genes which were 2-fold up or downregulated in either HE2 or progenitors following induction of each RUNX1 mutant. Left table shows the number of genes which were mutually up (red) or downregulated (blue), right table shows the number of genes which were upregulated in the dataset shown along the top and downregulated in the dataset on the side. Columns or rows which are greyed out have 0 genes deregulated in one of the datasets therefore cannot have any in common. C Heatmap showing the Pearson of correlation with hierarchical clustering of the +dox/-dox fold change for all deregulated genes across all 8 datasets. D The percentage of up or downregulated genes associated with RUNX1, RUNX1-ETO or RUNX1-EVI1 ChIP peaks is plotted.

We then examined whether the mutant RUNX1 proteins were targeting the same transcriptional networks. We first performed pair-wise analysis, to see whether different mutant proteins cause different or similar changes in gene expression patterns (Figure 4B). This analysis showed that just under a quarter of the genes which were upregulated in HE2 after R204X induction were also upregulated in HE2 by RUNX1-EVI1; a greater number of these genes were downregulated in progenitors by RUNX1-ETO rather than upregulated indicating complex stage-specific regulation. In a similar vein, multiple genes which were up-regulated by RUNX1-EVI1 were both up or downregulated by RUNX1-ETO, in both HE2 and progenitors, and those genes which were down-regulated by RUNX1-EVI1 were predominately up-regulated by RUNX1-ETO particularly in progenitors indicating opposing regulatory mechanisms.

We then performed correlation analysis and hierarchical clustering based on all genes which were changing across all of the datasets (Figure 4C). This analysis showed the genes affected by RUNX1-EVI1 were mostly unique and to a lesser extent this was true for all the other mutant RUNX1 driven gene expression changes indicating a mutant-specific pattern of gene expression changes. However, this analysis also indicated an inverse correlation in gene expression changes caused by R204X and RUNX1-ETO which was of note as these both drive AML and contain the RUNT-domain portion of RUNX1.

Next we analysed which of the genes with altered expression were direct targets of either RUNX1 or the two fusion proteins. None of the 3 genes deregulated by R201Q were RUNX targets. RUNX1-EVI1 caused genes to be down-regulated where it bound but most changes in gene expression were driven by the large scale increase in RUNX1 binding (Figure 4D, Figure 2) whereas loss of RUNX1, and RUNX1-ETO binding itself correlated with gene expression changes seen in response to RUNX1-ETO induction. A large proportion of the genes upregulated in response to R204X were RUNX1 targets but binding of RUNX1 was unchanged again indicating that this oncoprotein perturbs the action of RUNX1 at its binding sites rather than disrupting binding itself.

The impact of the RUNX1 oncoproteins on gene expression was therefore mild, varied and generally occurred in a mutation specific fashion, despite a significant proportion of affected genes being RUNX1 targets.

### Mutant oncoproteins disrupt RUNX1-mediated transcription factor and chromatin reorganisation

We previously showed that the up-regulation of RUNX1 during hematopoietic specification leads to a global reorganisation of transcription factors binding and chromatin patterns (Gilmour *et al*, 2018; Lichtinger *et al*, 2012). We therefore hypothesised that the RUNX1 mutants may interfere with this process and disrupt the transcription factor hubs that provide instruction for further blood cell differentiation. We first examined the transcription factor binding motifs associated with differential chromatin accessibility and found that the patterns of motif enrichment were specific to each RUNX1 mutant (Figure 5A, Supplementary Figure 5A). With R201Q we found an increase in chromatin accessibility associated with GATA motifs, with RUNX1-ETO accessible sites associated with RUNX and PU.1 were lost, and with RUNX1-EVI1 sites containing GATA and RUNX motifs were lost but PU.1 sites were gained. Interestingly, following induction of R204X - which lacks a transactivation domain - accessible chromatin sites were both lost and gained (Figure 5B) but were not associated with any changes in motif enrichment. RUNX motifs were also unchanged with R204X suggesting it is not acting dominant negative to the endogenous RUNX1 which again echoes the phenotypic and ChIP-seq data.

**Figure 5 -.**
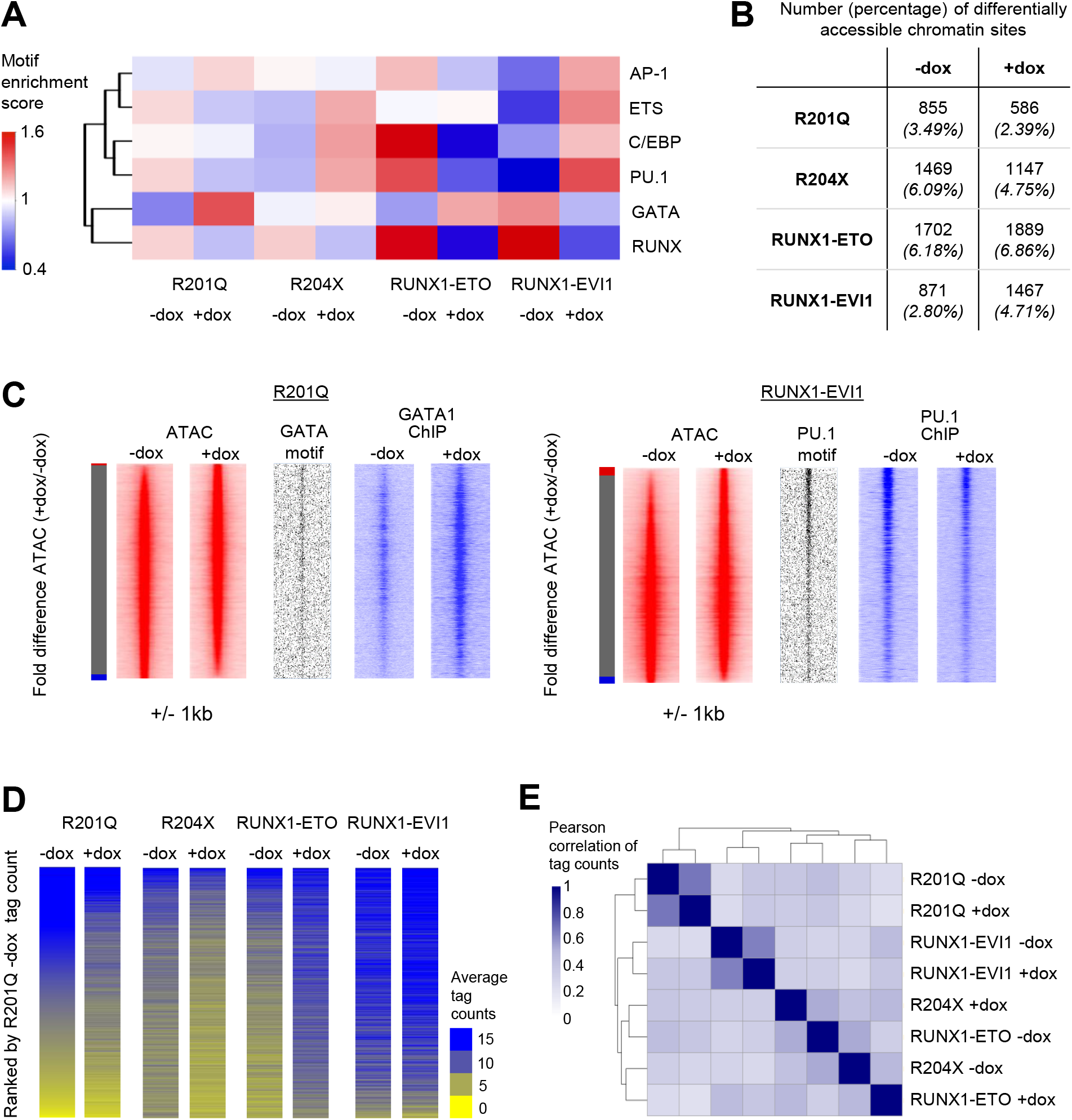
Chromatin accessibility changes are unique to each RUNX1 mutant and correlate with specific transcription factor binding patterns. A Heatmap with hierarchical clustering, showing the normalised enrichment score for transcription factor motifs which were seen in the de novo motif search of specific distal ATAC sites. B Table showing the number of specific ATAC peaks, and the percentage of the total peaks this corresponds to. C Chromatin accessibility in progenitors was ranked by fold change of the +dox/-dox tag count and represented as density plots (+/- 1kb). Motif enrichment and ChIP-seq of key transcription factors are plotted alongside. D ATAC tag counts were calculated across union of all -/+dox specific distal peaks across all four RUNX1 inductions in progenitors, and ranked according to R201Q -dox descending tag count. 9494 unique peaks out of 9986 used were unique. E Heatmap showing the Pearson correlation and hierarchical clustering which was performed using the tag counts of the union of specific peaks calculated in (B).

We confirmed two of these changes in motif composition by performing ChIP-seq for the transcription factors which bind to them. With R201Q, increased accessibility at GATA motifs was associated with overall increased GATA1 (the GATA factor most highly expressed in progenitors) binding; whereas with RUNX1-EVI1, PU.1 binding was maintained but was more prevalent at those sites where chromatin accessibility was gained (Figure 5C). These results highlight a profound disturbance of RUNX1-driven transcription factor binding reorganisation.

Alongside the changes associated with transcription factor binding we investigated whether lost or gained ATAC-seq peaks were shared or specific for each RUNX1 mutant. All –dox samples were generally well correlated allowing a comparison between changes caused by each oncoprotein (Supplementary Figure 5B). We calculated the union of all differential peaks and ranked them in parallel, ordered by the R201Q -dox sample (Figure 5D). As with the RNA-seq experiments (Figure 4C), this analysis again showed that each mutant RUNX1 changed the accessible chromatin landscape in a specific fashion with only a few common differentially accessible regions. We noted an inverse pattern of changes caused by R204X and RUNX1-ETO. To further examine this finding, we performed a correlation analysis using the tag counts for each sample across all differentially accessible peaks (Figure 5E), which again showed that R204X +dox and RUNX1-ETO -dox and R204X -dox and RUNX1-ETO +dox each cluster together although the majority of differentially accessible peaks were still unique (Supplementary Figure 5C). Taken together with the RNA-seq, these data suggest that R204X and RUNX1-ETO induction affected similar networks but not necessarily in the same way and this may be why they cause a similar phenotypic outcome.

RUNX1 transcription factor complexes can include histone acetyltransferases which RUNX1-ETO in particular is known to disrupt (Amann *et al.*, 2001; Wang *et al*, 1998). We therefore examined whether RUNX1-mutant specific chromatin changes were associated with altered histone acetylation patterns. Global H3K27ac patterns were dramatically affected by R204X, RUNX1-ETO and RUNX1-EVI1 induction, with acetylation both lost and gained around accessible chromatin (Figure 6A).

**Figure 6 -.**
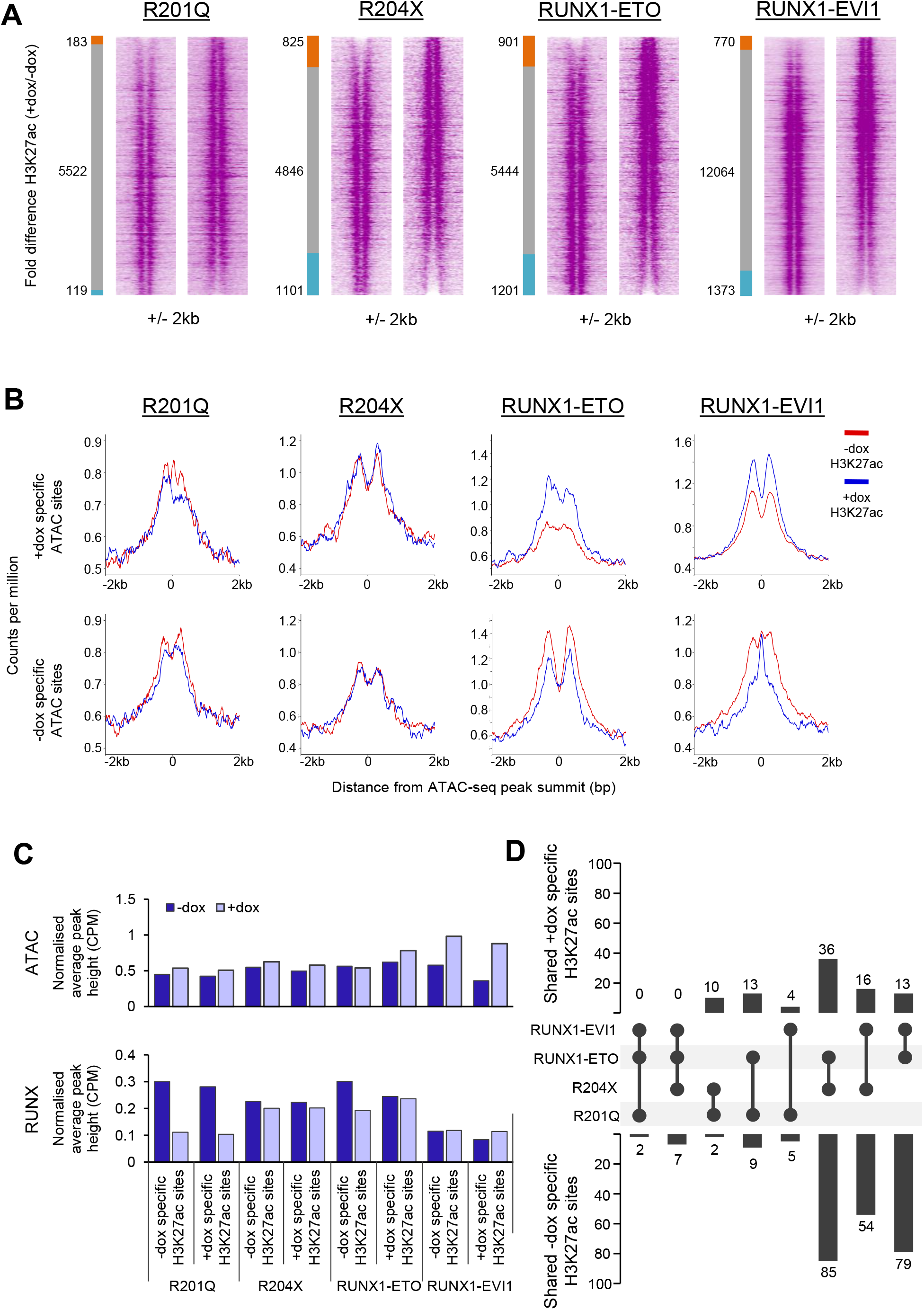
H3K27ac changes caused by RUNX1 mutants are not wholly dependent on changing chromatin accessibility. A The H3K27ac ChIP-seq signal at open chromatin sites in progenitors was ranked by fold change of the +dox/-dox tag count and represented as density plots (+/- 2kb). The side bar indicates +dox specific sites (orange), grey shared and blue -dox specific sites where specific sites are at least 2-fold different. The number of sites is indicated. B Average profiles of H3K27ac CPM-normalised ChIP-seq signal in progenitors plotted around the differential distal ATAC sites identified in Fig 5 (+/- 2kb). C The CPM-normalised average peak heights of ATAC-seq and RUNX1 ChIP-seq were calculated for the specific sites identified in (B). D The percentage of shared specific sites identified in (B) was calculated and shown by the bar graphs, where the circles indicate sets which have been overlapped in each case. Sets where there are no intersecting sites in either the - or +dox specific sites are not shown.

Lost or gained histone acetylation was not exclusively linked to lost or gained chromatin accessibility. The small number of chromatin changes observed in response to R201Q or R204X expression were reflected in the H3K27ac alterations at these sites (Figure 6B, Supplementary Figure 6A), but levels of H3K27ac strongly increased or decreased at sites with similarly altered chromatin accessibility following RUNX1-ETO and RUNX1-EVI1 binding, coinciding with up or downregulation of the associated genes (Figure 2A). Interestingly, at accessible chromatin sites lost after RUNX1-EVI1 expression (Figure 6B), we observed a pattern consistent with the flanking histones moving together indicative of a loss of transcription factor complexes at these sites (Bevington *et al*, 2017).

As expected from the initial analysis, the differential H3K27ac sites were not associated with differential chromatin accessibility except in the case of RUNX1-ETO (Figure 6C, Supplementary Figure 6B). Furthermore, differential H3K27ac sites were only minimally linked to altered RUNX1 binding, again predominately following RUNX1-ETO induction, suggesting whilst these changes are occurring where the RUNX1 complexes bind, they result from perturbation of the larger complex rather than just defects in RUNX1 binding. We also noted that a greater proportion of the sites which lost H3K27ac following induction of R204X, RUNX1-ETO or RUNX1-EVI1 were shared as compared to the sites which gained H3K27ac (Figure 6D), which may indicate these proteins are acting more similarly as repressors here.

Collectively, our data show that in spite of the relatively modest changes in gene expression, the induction of all RUNX1 oncoproteins interferes with RUNX1 activity and rapidly alters transcription factor occupancy and histone modification patterns.

### Mutant RUNX1 proteins alter lineage-specific chromatin priming

The developmentally controlled activation of differential gene expression programs during haematopoietic specification requires the gradual reorganization of chromatin preceding the onset of tissue specific gene expression, known as chromatin priming (Bonifer & Cockerill, 2017; Goode *et al*, 2016). Since RUNX1 is essential for the establishment of a blood-cell specific chromatin landscape (Lichtinger *et al*, 2012), we hypothesised that despite causing minimal alterations in gene expression, each mutant RUNX1 protein may uniquely perturb the chromatin architecture and transcription factor regulatory networks to differentially prime haematopoietic progenitor cells and thus derail future development.

We therefore analysed the degree to which the differentially accessible chromatin sites previously identified (Figure 2A) were shared with different precursor and mature cell types. The ATAC-seq data used for this analysis were derived from purified common myeloid progenitor (CMP), B-cell, Monocyte, Erythroblast and Megakaryocyte in order to cover the key lineage branches which RUNX1 mutation is known to influence (Heuston *et al*, 2018; Lara-Astiaso *et al*, 2014). A cell-type specific chromatin signature was calculated for each cell-type by identifying only those peaks which were not shared between different cell types. This set of peaks was compared to the differentially accessible chromatin sites formed or lost after induction of RUNX1 mutant proteins, as shown in the schematic in Figure 7A. An enrichment (Z) score was determined by comparing them to randomly sampled peaks within the union of all accessible sites for all cell types. In a healthy progenitor cell, we would expect to see balanced lineage priming for mature cells, as well as the progenitor cell signature. By examining the specifically lost or gained sites we could therefore understand how the RUNX1 mutants perturbed lineage priming.

**Figure 7 -.**
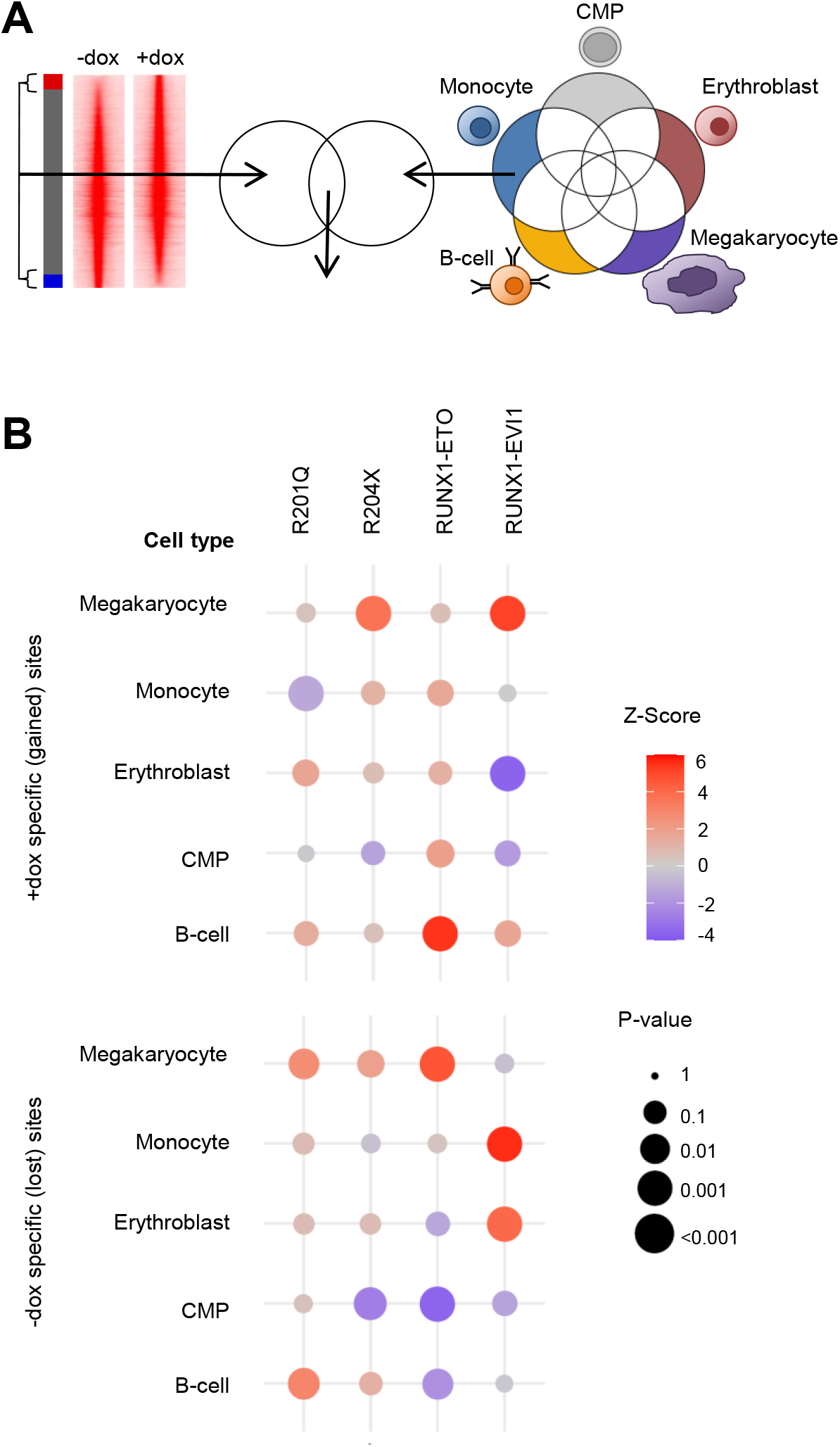
RUNX1 mutants disrupt RUNX1 driven chromatin priming. A Scheme of how the enrichment of differentially accessible ATAC-seq peaks from Figure 2A, intersecting with ATAC peaks specific to CMPs, B-cells, monocytes, erythroblasts or megakaryocytes was calculated. B Bubble plots showing the association of differentially accessible peaks after mutant RUNX1 induction with each peak set from the indicated cell types. Each bubble represents one intersection, the Z-score representing level of enrichment (red) or depletion of sites of each lineage (blue) as shown by the colour scale. The p-value is shown by the size of the circle.

After R201Q induction, we found that lost accessible chromatin sites (Figure 7B, lower panel) were highly enriched for a megakaryocyte signature, whereas those sites which were gained showed a slight enrichment for the erythroid fate (Figure 7B, upper panel), indicative of a skew in the megakaryocyte-erythroid branch of blood cell development (Figure 7B, Supplementary Figure 7). Both lost and gained sites also showed enrichment for B-cell primed sites, although to a lesser degree in the gained sites. None of the sites which gained chromatin accessibility were associated specifically with the monocyte lineage. A different pattern of changes was seen with R204X induction, where megakaryocyte priming was strongly enriched in sites where chromatin accessibility was gained, but also slightly enriched in lost sites as well, again suggesting a disruption of differentiation rather than a clear change of cell fate. Priming for all other lineages was preserved, but we found a significant absence of sites associated with CMPs in sites which lost accessibility following expression of R204X suggesting a preservation of the CMP chromatin state.

Both the fusion oncoproteins caused a greater disruption in the balance of lineage priming, in line with them causing increased phenotypic and gene expression changes. RUNX1-ETO led to a gain in accessibility at sites associated with both CMPs and the B-cell lineage, an example of which is shown in Supplementary Figure 7, and with a reciprocal lack of these lineages losing chromatin accessibility. At the same time, sites specific for the megakaryocyte lineage were lost, and a small proportion of sites associated with the monocytic lineage were gained. Similarly, RUNX1-EVI1 cause widespread disruption of priming but with no one lineage specifically gained or lost. Following induction of RUNX1-EVI1, sites associated with B-cells and megakaryocytes were gained (which can also be seen in Supplementary Figure 7), and sites associated with monocytes and erythroblasts were lost. Concordantly, erythroblast lineage chromatin sites were not gained in response to RUNX1-EVI1, nor were CMP sites. In summary, our data show that RUNX1 mutant proteins each influence the RUNX1 driven reorganisation of chromatin accessibility and lineage priming in unique ways leading to a disturbance of differentiation trajectories

## Discussion

In this study, we show that mutations in *Runx1* give rise to proteins which uniquely disrupt the gene regulatory networks at the onset of blood cell differentiation. During EHT, RUNX1 reorganises the transcriptional machinery to repress the endothelial fate and primes chromatin for continued hematopoietic differentiation (Gilmour *et al*, 2018; Lie-A-Ling *et al*, 2014). Chromatin priming at this stage by RUNX1 binding and elevated histone acetylation is critical for the correct binding patterns of transcription factors driving differentiation, such as PU.1 (Creyghton *et al*, 2010; Lichtinger *et al*, 2012). Here, we see a profound impact on chromatin priming as a result of perturbation of RUNX1 function at this stage. Most importantly, for RUNX1 point mutants, this perturbation occurred with only minimal influence on gene expression. RUNX1-ETO affected chromatin accessibility associated with RUNX, PU.1 and C/EBP motifs, leading to a skew in the progenitor/myeloid path as well as B-cell lineage, as has been previously implicated in t(8;21) leukaemia (Pabst *et al*, 2001; Ray *et al*, 2013; Sun *et al*, 2013; Tagoh *et al*, 2006). Conversely, R201Q caused gain of GATA1 binding. It was previously hypothesised that impaired erythropoiesis caused by RUNX1-DBD mutants was due to a change in RUNX1/GATA1 balance at the onset of erythroid differentiation (Cammenga *et al*, 2007; Waltzer *et al*, 2003). Our global binding data confirm this idea. RUNX1 is normally required to block the erythroid fate in favour of the megakaryocyte fate (Kuvardina *et al*, 2015; Song *et al*, 1999). Megakaryocytic differentiation is therefore dependent on the RUNX1/GATA1 balance as well (Elagib *et al*, 2003), suggesting a likely mechanism by which these RUNX1-DBD mutants contribute to platelet disorders.

One outstanding question has been the degree to which RUNX1-mutant phenotypes result from haploinsufficiency of RUNX1 due to the mutant proteins being non-functional or acting in a dominant negative fashion. Previous studies expressing mutant RUNX1 proteins in mice have shown them to have weakly dominant negative or null activity whereby blood cell formation was inhibited (Matheny *et al*, 2007). Some aspects of the mechanism by which the phenotype occurs can be inferred in these type of studies, such as inhibition of RUNX1-controlled myeloid gene expression (Guo *et al*, 2012) but due to the strong disruption in blood cell formation, the earliest events of cellular reprogramming by mutant RUNX1 proteins could not be studied. By inducibly expressing the mutant proteins on a background of wild type RUNX1 we demonstrate that all mutant proteins have additional functions.

From our data, we have developed a model of how these mutant RUNX1 proteins interact with the wild type RUNX1 to disrupt control of differentiation (Figure 8). Expression of the R201Q (DBD mutant) leads to a reduced interaction of wild type RUNX1 with CBFβ, a drastic reduction of global RUNX1 binding, increased GATA binding and thus a bias away from megakaryocyte differentiation. R204X (which lacks its TAD) does not affect wild type RUNX1 or other transcription factor binding but instead leads to changes in histone acetylation affecting the CMP trajectory. RUNX1-ETO displaces wild type RUNX1, leads to reduced expression and binding of PU.1 and C/EBPα (Pabst *et al*, 2001), and to reduced histone acetylation - this blocks cell differentiation at the early multipotent precursor cell stage and primes them towards a B-cell identity. RUNX1-EVI1 acts in a similar fashion to RUNX1-ETO but also causes increased RUNX1 binding associated with increased CBFβ interaction which has a knock-on effect on transcription factors such as PU.1 causing widespread disruption of all lineages in which RUNX1 is involved.

**Figure 8 -.**
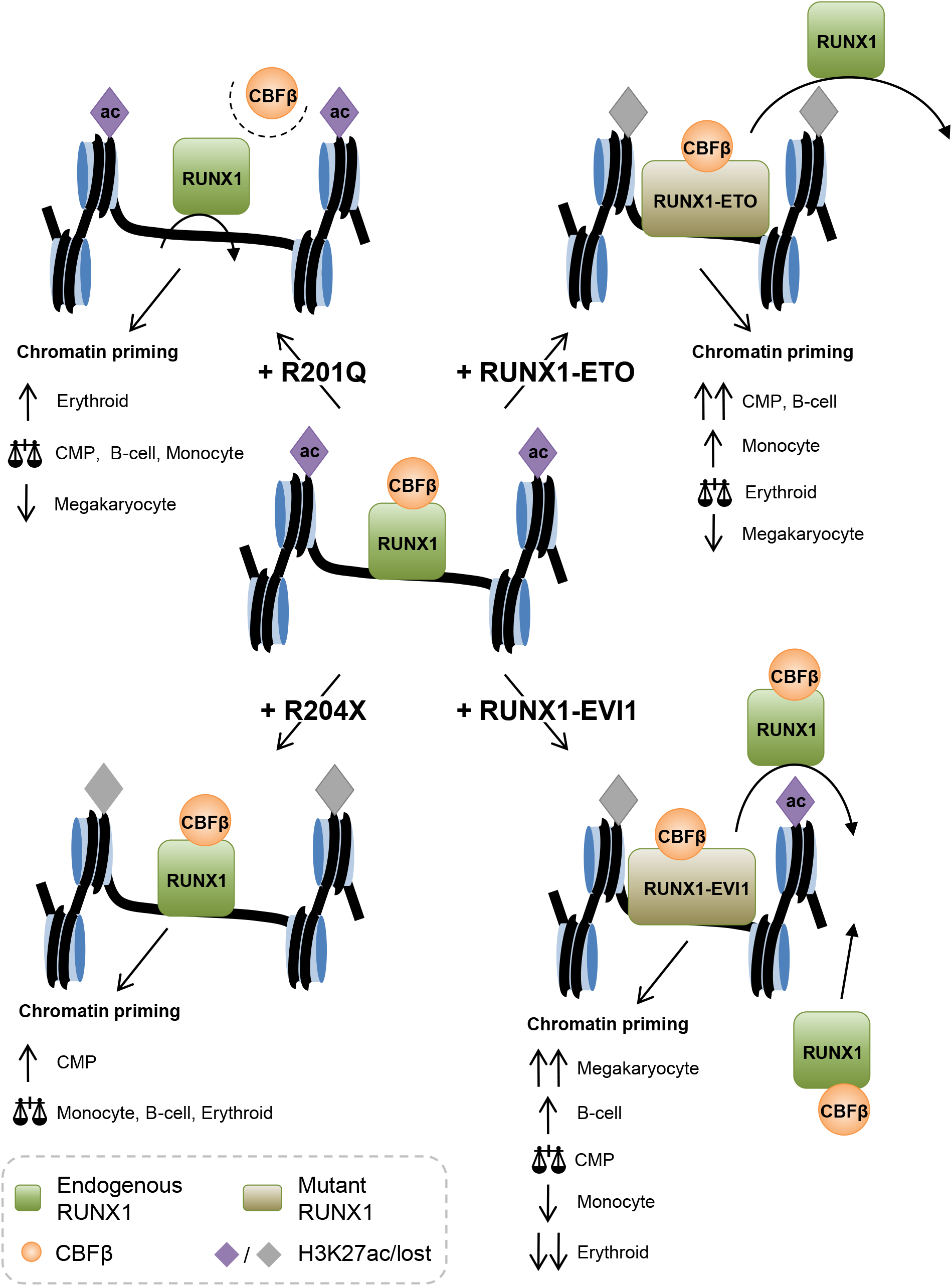
Mechanism of RUNX1 oncoprotein action on chromatin priming. A model for how each of the RUNX1 mutants is disrupting the normal activity of wild type RUNX1 (centre) based on the data we have generated.

In summary, our study has elucidated the genome-wide changes caused by four mutant RUNX1 proteins and shown that they disrupt the earliest instructions for the differentiation trajectory of haematopoietic progenitors. Full-length RUNX1 is required to rescue haematopoiesis in *RUNX1* knockout embryos and to set up balanced haematopoiesis (Goyama *et al*, 2004), which will require the establishment of correct lineage priming at the chromatin level. RUNX1 mutant proteins that miss different domains of the protein disturb this process. The expression of RUNX1-ETO and RUNX1-EVI1 is incompatible with normal blood cell development (Maki *et al*, 2005; Yergeau *et al*, 1997). However, RUNX1 point mutations can run in families (Song *et al*, 1999) and permit haematopoiesis which is in line with the results shown here. Our data would therefore predict that affected individuals in such families would already display signs of deregulation in the chromatin patterns of their progenitor cells. Our results demonstrate that different classes of mutation in RUNX1 have unique multi-factorial mechanisms of contributing to disease and so development of novel treatments will require an individual approach.

## Methods

### Mouse RUNX1-EVI1 ESC generation

Generation of RUNX1-ETO and RUNX1-EVI1 containing ESCs was previously described (Kellaway *et al*, 2020; Regha *et al*, 2015). R201Q and R204X plasmids were generated by site-directed mutagenesis on wild type human *RUNX1c*, and N-terminal HA tags added using the following primers: R201Q forward 5’-CAGTGGATGGGCCCCAAGAACCTCGAAGAC-3’, reverse 5’-GTCTTCGAGGTTCTTGGGGCCCATCCACTG-3’ and R204X forward 5’-CCCCCTCGAGCCACCATG-3’, reverse 5’-GCCGATGATATCTCAAGGTTCTCG-3’. A2lox ESCs (a gift from Michael Kyba) were transduced with 20 μg of each plasmid using the 4D-Nucleofector (Lonza), mouse ES program and P3 primary cell kit. Note, R201Q, R204X and RUNX1-ETO all included N-terminal HA-tags.

Individual colonies were expanded and maintained on mouse embryonic feeder cells in ES cell medium, comprising DMEM (Sigma D6546), 15% FCS (Sigma ES-009), 100 units/ml penicillin, 100 μg/ml streptomycin, 1 mM sodium pyruvate, 1mM L-glutamine, 0.15 mM monothioglycerol, 1x non-essential amino acids and 10^3^ U/ml leukaemia inhibitory factor (ESGRO, Millipore) following 7 days of 300 μg/ml neomycin selection.

### ESC Differentiation

ESCs were differentiated as previously described (Gilmour *et al*, 2014; Regha *et al*, 2015) with the following modifications. FLK1+ cells were purified by magnetic cells sorting, using biotin-conjugated CD309 antibody (eBioscience), anti-biotin microbeads (Miltenyi Biotec) and LS columns (Miltenyi Biotec) following culture of embryoid bodies for between 3.25 and 3.75 days (cell line dependent). These FLK1+ cells were then cultured in gelatin-coated flasks – 1.2-1.4×10^6^ cells in a T150 flask to form the blast culture. After 1-2 days (cell line dependent), 0.1-0.5 μg/ml doxycycline was added as appropriate and cells were cultured in the same media for a further 18 hours prior to sorting for HE and progenitors.

### FACS

Cell populations were identified and sorted on day 2-3 of blast culture based on surface markers. Cells were stained with cKit-APC (BD pharmingen), Tie2-PE (eBioscience) and CD41-PE-Cy7 (eBioscience) and analyzed on a Cyan ADP flow cytometer (Beckman Coulter) with data analysis using FlowJo, or sorted on a FACS Aria cell sorter (BD Biosciences).

### CFU assays

Unsorted floating cells were taken from d2-3 of the blast culture and 5×10^3^ cells were seeded in 1 ml MethoCult (M3434 STEMCELL Technologies) per dish, in duplicate and counted after 10 days.

### Western blotting

20 μg of protein extracts in Laemmli buffer were run on a 4-20% gradient pre-cast gel (Bio-Rad) and transferred to nitrocellulose using Turbo transfer packs (Bio-Rad). Membranes were blocked using 5% milk in TBS-T, then RUNX1 (C-terminal: ab23980, Abcam, 1:3000 or N-terminal: sc-8563 N-20, Santa Cruz Biotechnology, 1:250) or anti-HA (H6908, Sigma, 1:1000) was applied overnight at 4°C in 5% milk in TBS-T. After washing in TBS-T, this was followed with HRP-conjugated anti-rabbit or anti-goat antibody (Cell Signalling Technologies), and enhanced chemiluminescent reagent (Amersham) applied and blot was visualised using a Gel Doc system (Bio-Rad). For loading controls membranes were stripped using Restore stripping buffer (Thermo Fisher Scientific) and GAPDH (ab8245, Abcam) was applied and visualised as above.

### RNA-seq

RNA was isolated from sorted cells using either the NucleoSpin RNA kit (Macherey-Nagel) or Trizol reagent (Thermo Fisher Scientific). RNA-seq libraries were prepared from two biological replicates using the True-Seq stranded total RNA kit (Illumina) and sequenced paired-end in a pool of 12 indexed libraries using a Next-Seq 500/550 high output kit v2 150 cycles (Illumina) at the Genomics Birmingham sequencing facility.

### ATAC-seq

ATAC-seq was performed essentially as described (Buenrostro *et al*, 2015), briefly, 50,000 cKit+CD41+Tie2-progenitors were sorted by FACS and transposed in 1x Tagment DNA buffer (Illumina), Tn5 transposase (Illumina) and 0.01% Digitonin (Promega) for 30 minutes at 37°C with agitation. For R204X, RUNX1-ETO and RUNX1-EVI1 samples the tagmentation buffer additionally contained 0.3x PBS and 0.1% Tween-20. DNA was purified using a minelute reaction clean up kit (Qiagen). DNA was amplified by PCR using Nextera primers and libraries were sequenced using a Next-Seq 500/550 high output kit v2 75 cycles (Illumina).

### ChIP-seq

ChIP was performed as previously described (Kellaway *et al*, 2020; Obier *et al*, 2016) with the following modifications. cKit+ progenitors were sorted by MACS, and for PU.1 and H3K27ac crosslinked only in 1% formaldehyde (single crosslinking), or with both 415 μg/ml DSG, followed by formaldehyde (double crosslinking) for RUNX1 and GATA1. For single crosslinked cells nuclei were sonicated for 4 cycles of 30s on/30s off using a Picoruptor (Diagenode). Immunoprecipitation was carried out overnight at 4°C using 2 μg of RUNX1 antibody (ab23980, Abcam), PU.1 antibody (sc-352, Santa Cruz) or GATA1 antibody (ab11852, Abcam), or for four hours at 4°C using 1 μg of H3K27ac antibody (ab4729, Abcam) coupled to 15 μg Dynabeads Protein G (Invitrogen) per 2 x 10^6^ cells. DNA from 2-3 immunoprecipitations was pooled for RUNX1, but just 1 immunoprecipitation for H3K27ac, and extracted using Ampure beads (Beckman Coulter). ChIP libraries were generated using the KAPA hyper prep kit, libraries were size selected to obtain fragments between 150 and 450 bp and were sequenced as for ATAC-seq.

### Immunocytochemistry

5×10^5^ cells were adhered to microscope slides using a Cytospin cytocentrifuge (Thermo Fisher Scientific) for 3 minutes at 800 rpm, and fixed in 4% formaldehyde (Pierce) for 15 minutes. Cells were permeabilised in 0.1% Triton X-100 and non-specific staining was prevented by incubation in 3% bovine serum albumin. Antibodies were applied for 1 hour at room temperature prior to washing, anti-HA (H6908, Sigma) at 1:200, anti-EVI1 (2593, Cell Signalling Technology) at 1:200, anti-RUNX1 (sc-28679 H-65, Santa Cruz Biotechnology) at 1:200 or anti-CBFβ (sc-56751, Santa Cruz Biotechnology) and secondary Alexa Fluor 488-conjugated anti-rabbit (Jackson ImmunoResearch) at 1:200. Slides were mounted with ProLong Gold antifade reagent with DAPI (Invitrogen). Slides were visualised using a Zeiss LSM 780 equipped with a Quasar spectral (GaAsP) detection system, using a Plan Achromat 40x 1.2NA water immersion objective, Lasos 30mW Diode 405nm, Lasos 25mW LGN30001 Argon 488 and Lasos 2mW HeNe 594nm laser lines. Images were acquired using Zen black version 2.1. Post-acquisition brightness and contrast adjustment was performed uniformly across the entire image.

### Proximity ligation assay

Cells were prepared, fixed and blocked as for immunocytochemistry. Primary antibodies (sources as for immunocytochemistry) were applied in pairs – anti-CBFβ at 1:100, with either anti-RUNX1 at 1:20, anti-HA at 1:250 or anti-EVI1 at 1:100 for 1 hour at room temperature. Probes, ligation and amplification solutions (Duolink, Sigma Aldrich) were then applied at 37°C according to the manufacturer’s instructions and slides were mounted in Duolink mounting medium with DAPI (Sigma Aldrich). Slides were visualised as for immunocytochemistry. Post-acquisition brightness and contrast adjustment was performed uniformly across the entire image.

### RNA-seq analysis

Raw paired-end reads were processed to remove low quality sequences with Trimmomatic v0.38 (Bolger *et al*, 2014). Processed reads were then aligned to the mouse genome (mm10) using Hisat2 v2.1.0 (Kim *et al*, 2015) with default parameters. Read counts were calculated using featureCounts v1.5.1 (Liao *et al*, 2013) with the options -p -s 2. Gene models from refSeq (O’Leary *et al*, 2015) were used as the reference transcriptome. Only genes that were detected with at least 50 reads in at least one sample were retained for further analysis. Differential gene expression analysis was carried out using the voom method (Law *et al*, 2014) in the limma package v3.40.6 (Ritchie *et al*, 2015) in R v3.6.1. A gene was considered to be differentially expressed if it had a fold-change of at least 2 and an adjusted p-value less than 0.05.

Clustering of gene expression data was carried out by first calculating pair-wise Pearson correlations of the log2-transformed fold changes for each pair of samples in R, these were then hierarchically clustered using complete linkage of the Euclidean distances and plotted as a heatmap in R.

### ATAC-seq analysis

Single-end reads from ATAC-seq experiments were processed with Trimmomatic v0.38. Reads were then aligned to the mouse genome (mm10) with Bowtie2 v2.2.3 (Langmead & Salzberg, 2012) using the options --very-sensitive-local. Potential PCR duplicates were removed from the alignments using the MarkDuplicates function in Picard v2.10.5 (http://broadinstitute.github.io/picard). Regions of open chromatin (peaks) were identified using MACS2 v2.1.1 (Zhang *et al*, 2008) using the options -B --trackline --nomodel. The resulting peaks were then filtered against the mm10 blacklist (Amemiya *et al*, 2019) to remove potential artefacts from the data. Peaks were then annotated as either promoter-proximal if within 1.5kb of a transcription start site (TSS), and as a distal element otherwise. Promoter-proximal and distal elements were treated separately in all further analysis.

To carry out differential chromatin accessibility analysis, a peak union was first constructed by merging peaks from the -dox and +dox samples that had summits within 400bp of each other using the merge function in bedtools v2.26.0 (Quinlan & Hall, 2010). A new summit position was then defined for these peaks as the mid-point between the original summits. The average tag-density in a 400bp window centered on the peak summits was retrieved from the bedGraph files produced by MACS2 during the peak calling step. This was done using the annotatePeaks.pl function in Homer v4.9.1 (Heinz *et al*, 2010) with the options - size 400 -bedGraph. These were then normalized as tags-per-million (TPM) in R v3.6.1 and further log2-transformed as log_2_(TPM + 1). A peak was considered to be differentially accessible if it had at least a 2 fold-difference between -dox and +dox conditions. A de-novo motif analysis was carried out in the sets of gained and lost peaks using the findMotifsGenome.pl function in Homer using the options -size 200 -noknown.

Hierarchical clustering of ATAC-seq data was carried out using the log2-transformed normalized tag-counts. Pairwise pearson correlation values were calculated for each pair of samples and clustered using complete linkage of the Euclidean distances and plotted as a heatmap in R.

Tag density plots were constructed by retrieving the tag-density in a 2kb window centered on the peak summits with the annotatePeaks.pl function in Homer with the options -size 2000 - hist 10 -ghist -bedGraph. These were then plotted as a heatmap using Java TreeView v1.1.6 (Saldanha, 2004).

### ChIP-seq analysis

RUNX1-ETO, RUNX1-EVI1 and the RUNX1 ChIP-seq datasets from the RUNX1-ETO and RUNX1-EVI1 expressing cells (Kellaway *et al*, 2020; Regha *et al*, 2015) were downloaded from the Gene Expression Omnibus (GEO) under accession numbers GSE64625 and GSE143460.

Sequencing reads from ChIP-seq experiments were processed, aligned and de-duplicated as described above for the ATAC-seq data. Peaks from the RUNX1, RUNX1-ETO, RUNX1-EVI1, GATA1 and PU.1 ChIP-seq data were called using MACS2 v2.6.1 with the options -- keep-dup all -B --trackline -q 0.01. Peaks from the H3K27ac ChIP-seq data were also called using MACS2, but with addition of the --broad option. Only peaks that were found within open chromatin, as measured by the ATAC-seq data were retained for further analysis. Differential peak analysis was carried out in the same way as the differential chromatin accessibility analysis described above for the ATAC-seq data with a modification for the H3K27ac data for which the window to calculate the tag-density was increased to 800bp in order to count reads which flank the open chromatin. To identify potential targets for each of the transcription factors measured, we annotated the peaks to their closest gene using the annotatePeaks.pl function in homer v4.9.1. Average profiles were constructed from the ChIP-seq data using deepTools v3.3.2 (Ramírez *et al*, 2016). To do this, read counts were calculated and normalized as counts per million (CPM) using the bamCoverage function in deepTools, the average profile calculated using the computeMatrix function with the reference-point option, and then plotted in R. Shared sites were calculated using bedtools intersect and plotted using the UpSetR function in R. Tag-density plots were constructed as described above for the ATAC-seq data.

### Motif enrichment analysis

To identify transcription factor binding motifs that are enriched in a set of peaks relative to another, we calculated a motif enrichment score, S_ij_ for each motif i in each peak set j as

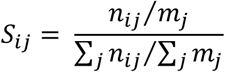

where n_ij_ is the number of sites in peak set j that contain the motif i, and mj is the total number of sites in peak set j. This was calculated for each TF motif in each of the peak sets being considered, and produced a matrix of enrichment scores which were then hierarchically clustered using complete linkage of the Euclidean distance in R and displayed as a heatmap. The set of motif probability weight matrices (PWMs) used for this analysis were derived from a de-novo motif search of the gained and lost ATAC-seq peaks using Homer.

Motif density plots were constructed by retrieving the motif density in a 2kb window centered on the peak summits with the annotatePeaks.pl function in homer with the options -size 2000 -hist 10 -ghist –m, using the Homer known motif database. These were then plotted as a heatmap using Java TreeView v1.1.6.

### Lineage priming analysis

In order to determine if the sets of +dox and -dox specific ATAC peaks that we found may also contain a chromatin signature that is normally only associated with a particular cell type, we carried out an analysis designed to measure if the number of cell type specific sites that are also found in our ATAC-seq data is significantly different than what would be expected by chance.

To do this we downloaded a set of ATAC-seq data that were generated from a number of mature cells types by (Heuston *et al*, 2018 and Lara-Astiaso *et al*, 2014). These data were downloaded from the Gene Expression Omnibus (GEO) under accession numbers GSE59992 and GSE143270. The cell types considered here were CMPs, B-cells, monocytes, erythroblasts and megakaryocytes. These ATAC-seq data were aligned and peaks were called and filtered as described above. Only peaks that were found in both replicates for each cell type were retained for further analysis. A peak was then considered to be cell type specific if it was found in only one of the cell types. This was done by comparing the peaks from each cell type to the union of peaks from all other cell types using the intersect function in bedtools with the -v parameter.

To determine if any of these cell type specific peak sets were either significantly enriched or depleted in our data, we carried out a randomisation based test for each of our RUNX1 mutant +dox and -dox specific peak sets as follows. First, we counted the number of +dox specific peaks that overlap with the cell type specific peaks using the intersect function in pybedtools (Dale *et al*, 2011). We then randomly sampled a set of peaks from the full set of distal sites in that RUNX1 mutant, and counted the number of overlapping peaks between this random set and the cell type specific peaks. The number of peaks sampled was equal to that of the +dox peaks, and could be sampled anywhere from the +dox, -dox and shared peaks. This procedure was repeated 1000 times and produced a list of counts measuring the overlap of the random sets with the cell type specific peaks. These counts were then used to calculate a Z-score using the formula

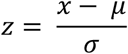

where x is the number of +dox peaks that overlap a cell type specific peak, μ is the mean of the counts from the re-sampling procedure, and σ is the standard deviation of those counts. A p-value measuring the statistical significance of the enrichment could also be derived from this test. This was calculated as the proportion of times the number of overlapping sites from the random peak sets was greater than that of the actual +dox peaks, with a low p-value suggesting that the number of cell type specific peaks found in the +dox peaks is greater than what would be expected only by chance. A p-value measuring depletion could also be calculated, and here is calculated as the proportion of times the number of overlapping sites from the random peak sets was less than that of the actual +dox peaks. In this case, a low p-value suggests that the cell type specific peaks are under-represented in the +dox specific peaks. This same test was also applied to each of the -dox specific peak sets.

## Data Availability

All sequencing data from this publication have been deposited to GEO and assigned the identifier GSE154623.

## Acknowledgements

This work was funded by grants from the Kay Kendall Leukaemia Fund (KKLF), the Biotechnology and Biological Sciences Research Council (BBSRC) and Blood Cancer UK (Bloodwise) to CB. We thank Genomics Birmingham for their expert sequencing service, the University of Birmingham Flow Cytometry unit for cell sorting, and Martin Higgs from the Institute of Cancer and Genomic Sciences for help with the PLA assay.

## Author Contributions

SGK designed and performed experiments and analysis, and wrote the manuscript. PK and BEW performed analysis. EK performed experiments, RK generated cell lines. CB conceived and coordinated the project, and wrote the manuscript. All authors commented on the manuscript.

