## Supplementary Figures for "Different mutant RUNX1 oncoproteins program alternate haematopoietic differentiation trajectories"

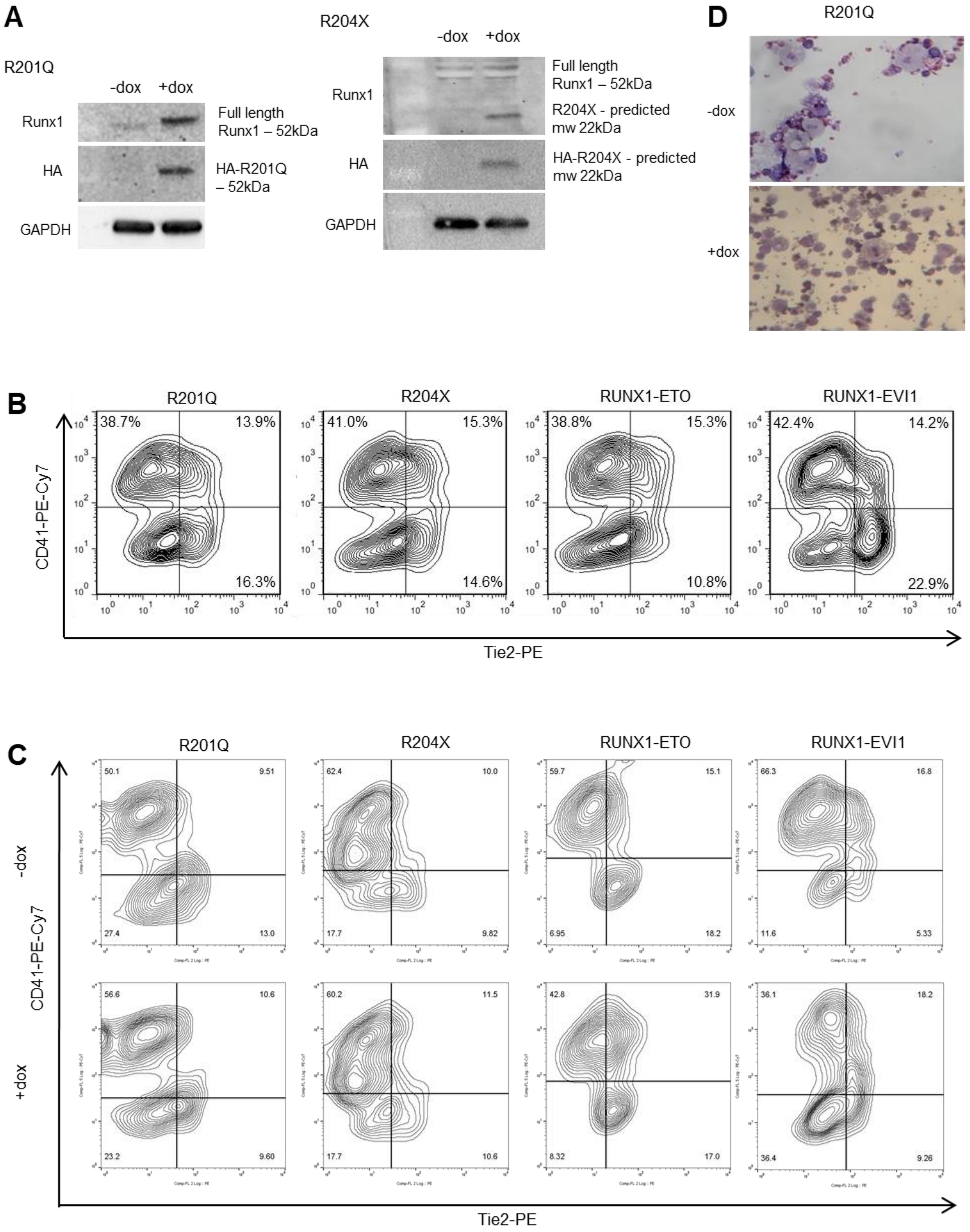
A
R201Q
-dox
+dox
Runx1
Full length
Runx1 – 52kDa
HA
HA-R201Q
– 52kDa
GAPDH
R204X
-dox
+dox
Runx1
Full length
Runx1 – 52kDa
R204X - predicted
mw 22kDa
HA
HA-R204X - predicted
mw 22kDa
GAPDH
D
R201Q
-dox
+dox
B
R201Q
R204X
RUNX1-ETO
RUNX1-EVI1
CD41-PE-Cy7
38.7%
13.9%
16.3%
41.0%
15.3%
14.6%
38.8%
15.3%
10.8%
42.4%
14.2%
22.9%
Tie2-PE
C
R201Q
R204X
RUNX1-ETO
RUNX1-EVI1
-dox
+dox
CD41-PE-Cy7
50.1
9.51
27.4
13.0
62.4
10.0
17.7
9.82
59.7
15.1
6.95
18.2
66.3
16.8
11.6
5.33
56.6
10.6
23.2
9.60
60.2
11.5
17.7
10.6
42.8
31.9
8.32
17.0
36.1
18.2
36.4
9.26
Tie2-PE

**Supplementary Figure 1 - Induction of RUNX1 mutants during blood differentiation perturbs progenitor identity**

A Mutant RUNX1 induction was not seen in the absence of dox shown by western blot using an antibody against the HA-tag, dox concentration required for induction was titrated such that expression was physiological as shown by the RUNX1 antibodies.

B Differentiation timings were adjusted such that induction of each mutant RUNX1 occurred at the same point of blast culture, in the same cell populations to minimise clonal variation, this was assessed using flow cytometry for Tie2-PE and CD41-PE-Cy7, pre-gated on cKit-APC, representative plots are shown from 3 biological replicates.

C  Representative flow cytometry plots from at least 3 biological replicates, pre-gated on cKit-APC are shown for the composition of the blast culture 18 hours following dox induction of the mutant RUNX1.

D Example images of Wright-Giemsa stained mixed lineage colonies from +/- R201Q.

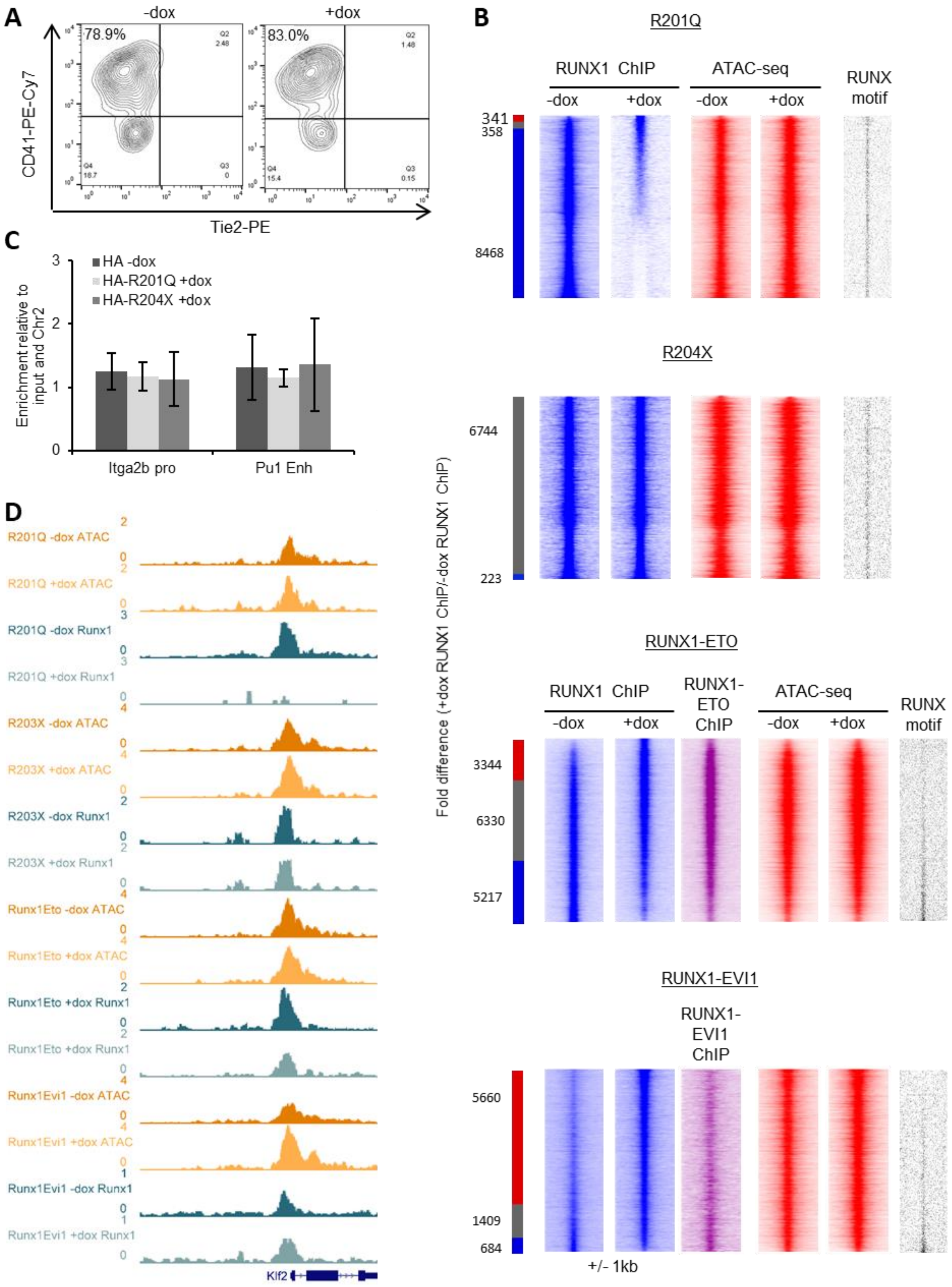
A
-dox
+dox
78.9%
83.0%
CD41-PE-Cy7
Tie2-PE
B
R201Q
RUNX1 ChIP
ATAC-seq
RUNX motif
-dox
+dox
341
358
8468
R204X
6744
223
RUNX1-ETO
RUNX1-ETO ChIP
3344
6330
5217
RUNX1-EVI1
RUNX1-EVI1 ChIP
5660
1409
684
+/- 1kb
Fold difference (+dox RUNX1 ChIP/-dox RUNX1 ChIP)
C
HA -dox
HA-R201Q +dox
HA-R204X +dox
Enrichment relative to input and Chr2
Itga2b pro
Pu1 Enh
D
R201Q -dox ATAC
R201Q +dox ATAC
R201Q -dox Runx1
R201Q +dox Runx1
R203X -dox ATAC
R203X +dox ATAC
R203X -dox Runx1
R203X +dox Runx1
Runx1Eto -dox ATAC
Runx1Eto +dox ATAC
Runx1Eto +dox Runx1
Runx1Eto +dox Runx1
Runx1Evi1 -dox ATAC
Runx1Evi1 +dox ATAC
Runx1Evi1 -dox Runx1
Runx1Evi1 +dox Runx1
Klf2

**Supplementary Figure 2 - Mutant RUNX1 induction leads to specific changes to endogenous RUNX1 binding and chromatin accessibility**

A Representative flow cytometry plots of the cKit-sorted progenitors used for ChIP experiments, shown here for R204X, pre-gated on cKit-APC.

B Endogenous RUNX1 ChIP-seq data in progenitors at both distal sites and promoters was ranked by fold change of the +dox/-dox tag count and represented as density plots. Chromatin accessibility and density of the RUNX1 motif are plotted alongside based upon the RUNX1 ranking. All density plots are centered around +/- 1kb of the RUNX1 ChIP peak summits. The red bar indicates +dox specific sites, grey shared and blue -dox specific sites where specific sites are at least 2-fold different. The numbers of sites are indicated.

C ChIP-qPCR was done using an antibody against the HA-tag following induction of R201Q and R204X, and in the absence of induction. The bars show the mean of 3 ChIP experiments, where enrichment of two RUNX1 positive control regions are compared to a negative control region, and to the input control. The error bars indicate standard error of the mean.

D UCSC Genome browser screenshot of CPM-normalised ATAC-seq and ChIP-seq tracks at the Klf2 promoter.

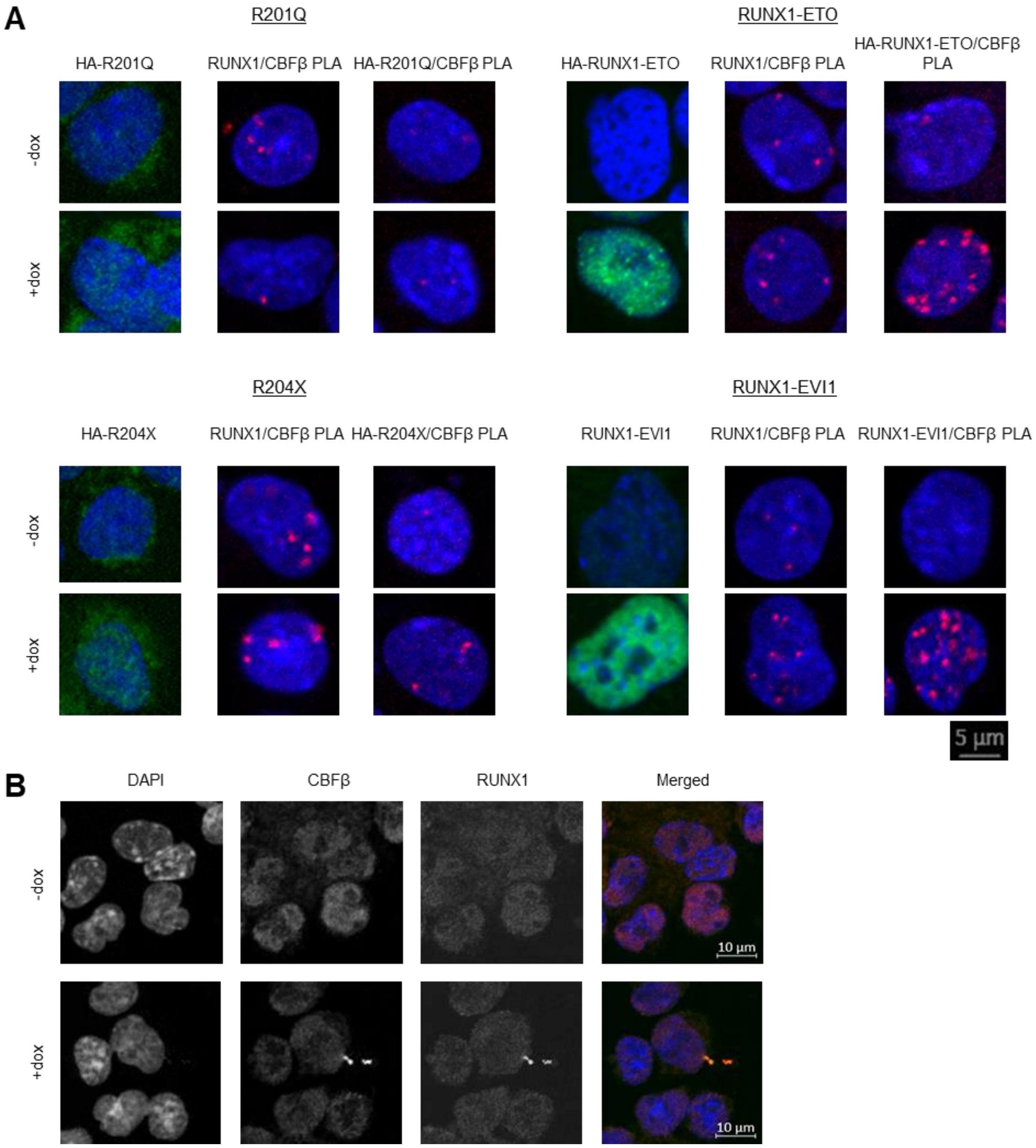
A
R201Q
RUNX1-ETO
HA-R201Q
RUNX1/CBFβ PLA
HA-R201Q/CBFβ PLA
HA-RUNX1-ETO
RUNX1/CBFβ PLA
HA-RUNX1-ETO/CBFβ PLA
-dox
+dox
R204X
RUNX1-EVI1
HA-R204X
RUNX1/CBFβ PLA
HA-R204X/CBFβ PLA
RUNX1-EVI1
RUNX1/CBFβ PLA
RUNX1-EVI1/CBFβ PLA
-dox
+dox
5 µm
B
DAPI
CBFβ
RUNX1
Merged
-dox
+dox
10 µm
10 µm

**Supplementary Figure 3 - RUNX1 mutants interact with CBFβ and partially disrupt RUNX1/CBFβ interactions**

A Enlarged images of the assays shown in Figure 3A. For each cell line, on the left is immunocytochemistry of the mutant protein alone (shown in green, using anti-HA or anti-EVI1 antibodies) counterstained with DAPI (blue). In the centre is PLA of endogenous RUNX1 with CBFβ (red), with DAPI (blue). On the right is PLA of the mutant RUNX1 with CBFβ (red), with DAPI (blue). Scale bar applies to all images.

B Immunocytochemistry controls for RUNX1 and CBFβ antibodies used in PLA assays shown here in R204X progenitors, individual channels are shown in grayscale with the merged pseudocolour image shown on the right, where DAPI is blue, CBFβ is red and RUNX1 is green.

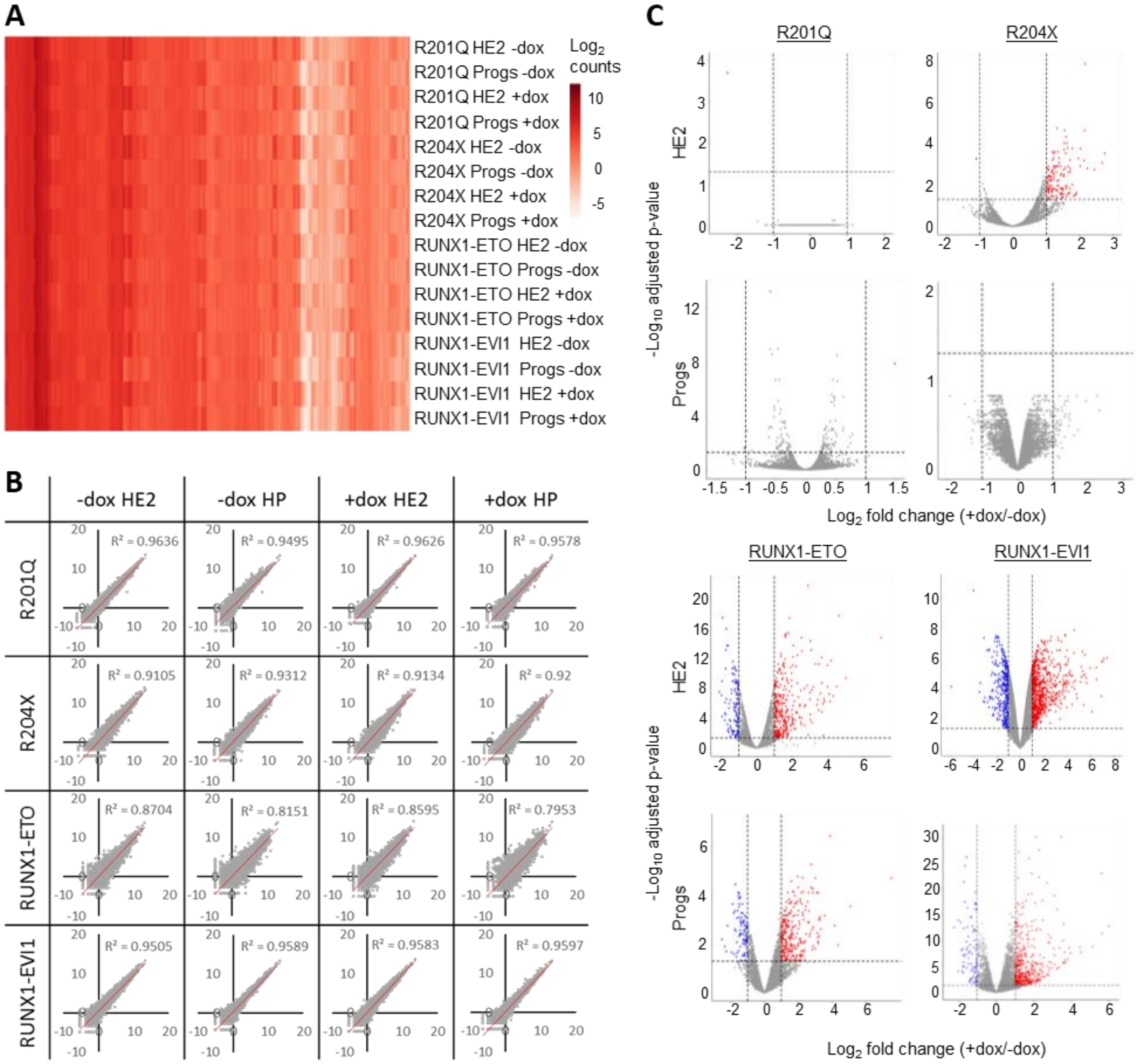

A
R201Q HE2 -dox
R201Q Progs -dox
R201Q HE2 +dox
R201Q Progs +dox
R204X HE2 -dox
R204X Progs -dox
R204X HE2 +dox
R204X Progs +dox
RUNX1-ETO HE2 -dox
RUNX1-ETO Progs -dox
RUNX1-ETO HE2 +dox
RUNX1-ETO Progs +dox
RUNX1-EVI1 HE2 -dox
RUNX1-EVI1 Progs -dox
RUNX1-EVI1 HE2 +dox
RUNX1-EVI1 Progs +dox
Log2 counts
B
-dox HE2
-dox HP
+dox HE2
+dox HP
R201Q
R204X
RUNX1-ETO
RUNX1-EVI1
R² = 0.9636
R² = 0.9495
R² = 0.9626
R² = 0.9578
R² = 0.9105
R² = 0.9312
R² = 0.9134
R² = 0.92
R² = 0.8704
R² = 0.8151
R² = 0.8595
R² = 0.7953
R² = 0.9505
R² = 0.9589
R² = 0.9583
R² = 0.9597
C
R201Q
R204X
RUNX1-ETO
RUNX1-EVI1
HE2
Progs
-Log10 adjusted p-value
Log2 fold change (+dox/-dox)

**Supplementary Figure 4 - Mutant forms of RUNX1 cause unique and shared gene expression changes**

A Heatmap showing the $log_2$ counts for all expressed genes for all RNA-seq samples used, following merging of duplicates.

B XY scatterplots of counts for all genes duplicates for all RNA-seq samples used, $R_2$ value is indicated based on best-fit line.

C Volcano plots of log2 fold change of +dox/-dox gene expression counts (x-axis) against p-value. Genes called as upregulated are indicated in red, and downregulated genes in blue, genes in grey were considered unchanged.

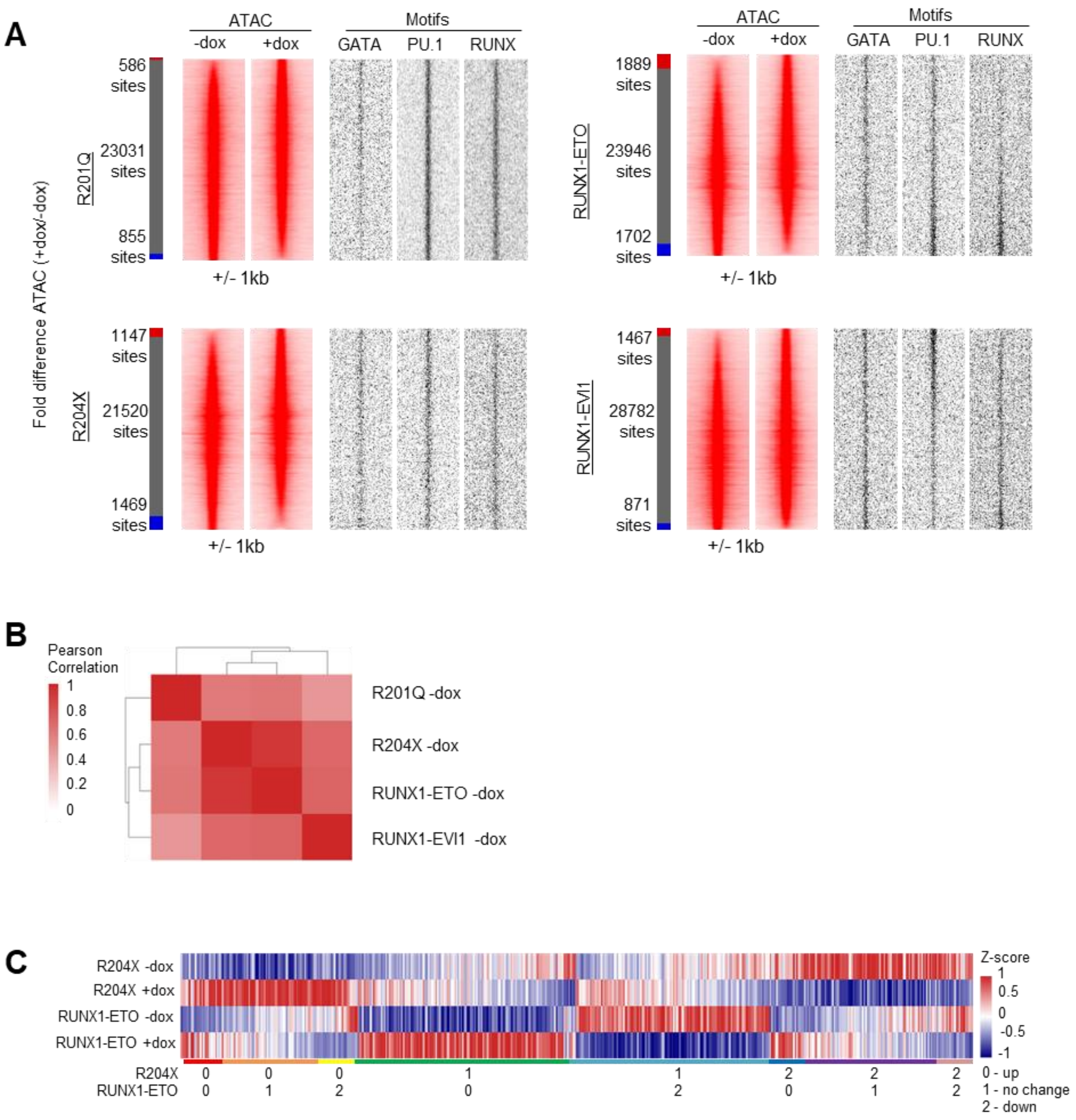
A
ATAC
-dox
+dox
Motifs
GATA
PU.1
RUNX
Fold difference ATAC (+dox/-dox)
R201Q
586 sites
23031 sites
855 sites
+/- 1kb
RUNX1-ETO
1889 sites
23946 sites
1702 sites
R204X
1147 sites
21520 sites
1469 sites
RUNX1-EVI1
1467 sites
28782 sites
871 sites
B
Pearson Correlation
1
0.8
0.6
0.4
0.2
0
R201Q -dox
R204X -dox
RUNX1-ETO -dox
RUNX1-EVI1 -dox
C
R204X -dox
R204X +dox
RUNX1-ETO -dox
RUNX1-ETO +dox
Z-score
1
0.5
0
-0.5
-1
R204X
RUNX1-ETO
0 - up
1 - no change
2 - down

**Supplementary Figure 5 - Chromatin accessibility changes are unique to each RUNX1 mutant and correlate with specific transcription factor binding patterns**

A Distal ATAC peaks were ordered according to the fold-difference of the normalised tag count between +dox and -dox progenitors and presented as a density plot for each sample (+/-1kb). Motif densities are plotted alongside for key transcription factors. The red bar indicates +dox specific sites, grey shared and blue -dox specific sites where specific sites are at least 2-fold different.

B Heatmap showing the Pearson correlation and hierarchical clustering of normalised tag counts for distal peaks in all uninduced ATAC-seq samples.

C Heatmap showing Z-score of average tag density at all peaks specifically gained or lost following induction of RUNX1-ETO or R204X. The coloured bar below indicates groups of whether genes are commonly up or down-regulated or unchanged.

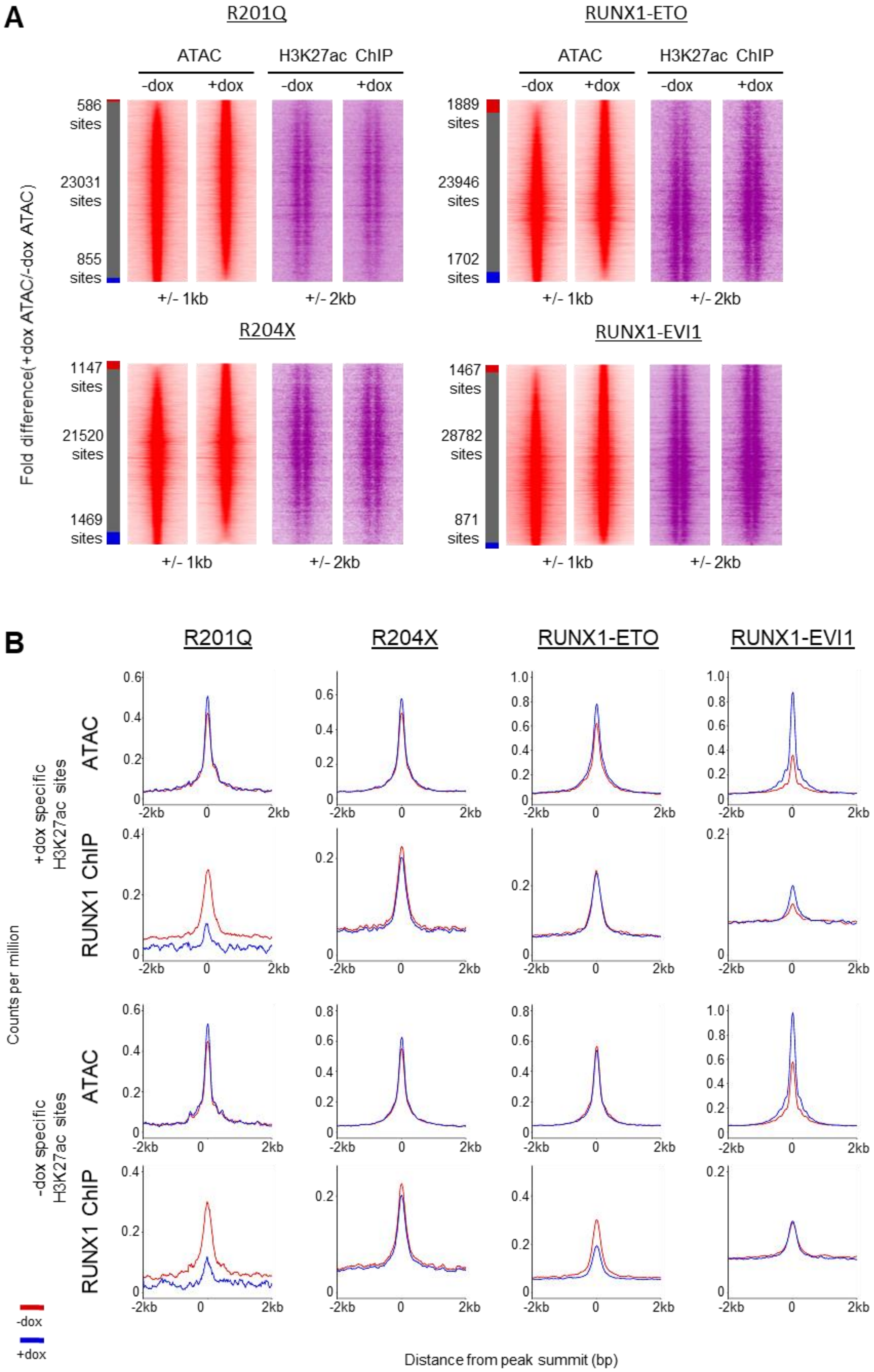
A
R201Q
RUNX1-ETO
ATAC
H3K27ac ChIP
-dox
+dox
586 sites
23031 sites
855 sites
1889 sites
23946 sites
1702 sites
+/- 1kb
+/- 2kb
R204X
RUNX1-EVI1
1147 sites
21520 sites
1469 sites
1467 sites
28782 sites
871 sites
Fold difference(+dox ATAC/-dox ATAC)
B
R201Q
R204X
RUNX1-ETO
RUNX1-EVI1
+dox specific H3K27ac sites
-dox specific H3K27ac sites
ATAC
RUNX1 ChIP
Counts per million
-2kb
0
2kb
-dox
+dox
Distance from peak summit (bp)

**Supplementary Figure 6 - H3K27ac changes caused by RUNX1 mutants are not wholly dependent on changing chromatin accessibility**

A Distal ATAC peaks were ordered according to the fold-difference of the normalised tag count between +dox and -dox progenitors and presented as a density plot for each sample (+/- 1kb). H3K27ac ChIP seq is plotted alongside (+/- 2kb from the ATAC peak summits).

B Average profiles of ATAC-seq and RUNX1 ChIP-seq signal around the H3K27ac specific sites calculated in Fig 6B (+/- 2kb from the ATAC peak summits).

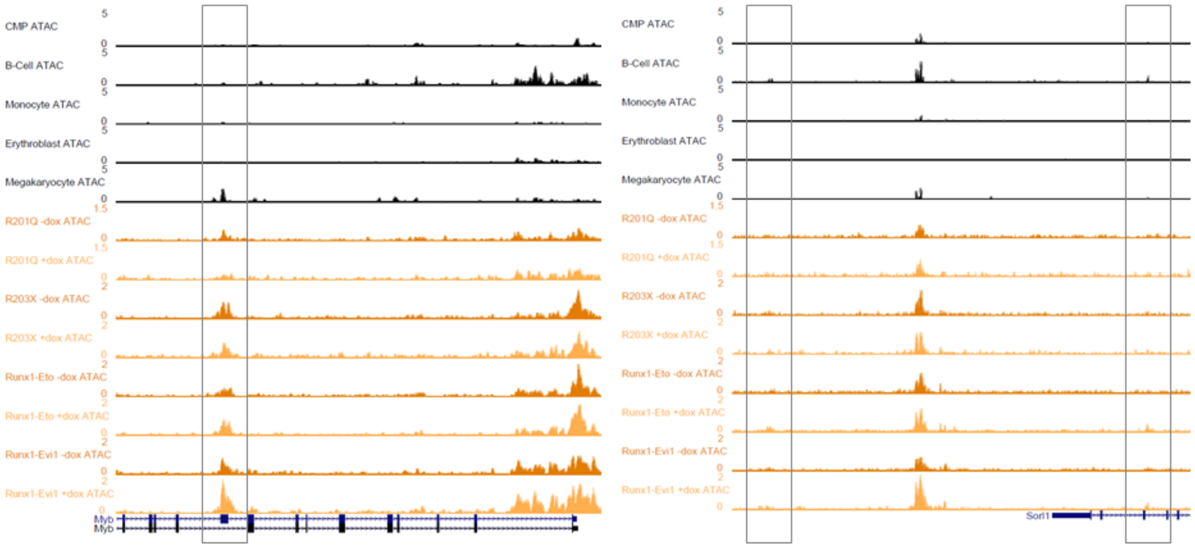


**Supplementary Figure 7 - RUNX1 mutants disrupt RUNX1 driven chromatin priming**

UCSC Genome Browser screenshots showing (left) a megakaryocye specific accessible chromatin site in the *Myb* gene which is lost with R201Q expression and gains accessibility with RUNX1-EVI1, and (right) two B-cell specific sites in the *Sorl1* gene which are gained with RUNX1-ETO.
